# Cellular energy sensor SnRK1 suppresses salicylic acid-dependent and -independent defenses and bacterial resistance in Arabidopsis

**DOI:** 10.1101/2025.10.01.679707

**Authors:** Linnan Jie, Miho Sanagi, Shigetaka Yasuda, Kohji Yamada, Saki Ejima, Ayumi Sugisaki, Junpei Takagi, Mika Nomoto, Xiu-Fang Xin, Yasuomi Tada, Yusuke Saijo, Takeo Sato

## Abstract

In nature, plants cope with various pathogens that compete for cellular resources during infection. It has long been suggested that plant defense activity must be linked to cellular energy and metabolic states to optimize the balance between growth and defense. However, the molecular mechanisms that regulate immune activity in relation to cellular energy status remain unclear. Here, we demonstrate that the plant energy sensor SNF1-RELATED KINASE 1 (SnRK1) plays a critical role in modulating defense responses and bacterial resistance in *Arabidopsis thaliana*. Bacterial elicitor-induced expression of defense marker genes, such as *PATHOGENESIS-RELATED 1* (*PR1*), is significantly repressed under sugar-limited conditions in wild-type seedlings, whereas this expression is markedly enhanced in the *snrk1* knockdown mutants. SnRK1 restricts defense-related gene expression and resistance to the biotrophic bacterial pathogen *Pseudomonas syringe* pv. *tomato* DC3000, which are partly dependent on salicylic acid (SA). In addition, we found that the SnRK1 kinase activity is increased by high humidity. Consistently, SnRK1 is critical for the suppression of SA-mediated defense responses under high humidity conditions. SnRK1 physically associates with the SA-related transcription factors TGACG SEQUENCE-SPECIFIC BINDING PROTEIN 4 (TGA4) and TGA2 to attenuate *PR1* expression. These findings provide valuable insight into the molecular mechanisms linking cellular energy status with immune regulation in plants.

## Introduction

Plants frequently encounter pathogen challenges in nature and have evolved a sophisticated innate immune system against microbial infection (1–3). Cell surface-localized pattern recognition receptors (PRRs) directly recognize microbe- or pathogen-associated molecular patterns (MAMPs/PAMPs), or endogenous damage-associated molecular patterns (DAMPs). This recognition triggers intracellular signal transduction cascades that activate defense responses, including transcriptional reprogramming and the production of defense-related phytohormones such as salicylic acid (SA), jasmonic acid and ethylene (4–6). SA is essential for resistance to a wide range of biotrophic and hemi-biotrophic pathogens (7–10). In *Arabidopsis thaliana* (hereafter Arabidopsis), SA biosynthesis is induced following pathogen attack mainly through the isochorismate (ICS) pathway (11, 12). SA is sensed by the proteins NONEXPRESSOR OF PATHOGEN-RELATED GENE 1 (NPR1) and NPR3/4, which act as defense-related transcriptional coregulators through associations with transcription factors, such as TGACG SEQUENCE-SPECIFIC BINDING PROTEINs (TGAs) (13, 14). These NPR-TGA modules directly regulate the expression of defense marker genes, such as *PATHOGENESIS-RELATED 1* (*PR1*) and *WRKY DNA-BINDING PROTEIN 38* (*WRKY38*), according to cellular SA levels in plants (15–25). Proteolytic processing of PR1 generates an immunomodulatory cytokine (CAPE9) to activate systemic immunity (26). *WRKY38* acts as negative regulator against SA-induced genes to fine-tune plant basal defense (27). It is becoming apparent that pathogen-induced SA biosynthesis and signaling are modulated by various environmental stimuli, thereby influencing plant susceptibility to pathogens (28–30).

Sugars serve as both essential nutrients and signaling molecules that optimize plant growth and metabolism (31–34). In Arabidopsis, glucose is sensed by HEXOKINASE 1 (HXK1), a dual functional protein acting as an enzyme and sensor in modulating global gene expression related to primary metabolism and growth (35). TARGET OF RAPAMYCIN (TOR) and SNF1-RELATED KINASE 1 (SnRK1) are master regulators that integrate nutrient availability status to fine-tune plant growth. Their phosphorylation target proteins include key enzymes and gene expression regulators related to metabolism and plant growth (36–38). In Arabidopsis, TOR promotes cell proliferation and protein synthesis under nutrient-replete conditions (39). In contrast, SnRK1 maintains cellular metabolism and energy homeostasis under energy-limited conditions, such as prolonged darkness and submergence, by inhibiting anabolic processes while promoting nutrient recycling and anaerobic metabolism (40–47). SnRK1 is the plant ortholog of the cellular energy sensors mammalian AMP-ACTIVATED KINASE (AMPK) and yeast SUCROSE NON-FERMENTING 1 (SNF1), and forms heterotrimeric complexes consisting of a catalytic α-subunit and regulatory β- and γ-subunits (48–50). In mammals and yeast, high glucose levels inhibit AMPK and SNF1 activities (51–54). Although the molecular basis of SnRK1 activity regulation is less clear compared with AMPK and SNF1, recent studies suggest that trehalose 6-phosphate (T6P) and class II T6P synthase (TPS)-like proteins directly interact with SnRK1, respectively, to negatively modulate its kinase activity (55, 56). SnRK1 also modulates developmental processes such as flowering and lateral root formation by phosphorylating key transcription factors involved in these processes (57, 58). Moreover, SnRK1 has been implicated in plant defense responses (59–61), although the underlying mechanisms remain poorly understood.

It has been suggested that plant immunity must be tightly coordinated with cellular energy and metabolic states to balance growth and defense (31, 62). Exogenous sugar application enhances pathogen resistance (63), whereas prolonged sugar starvation under darkness increases susceptibility (64). Recent research also indicates that glucose-6-phosphate plays a critical role in regulating defense-related transcriptional activation and SA production (65). Together, these findings suggest that sugars serve as critical integrators of energy status and defense regulation in plants. However, the molecular mechanisms that connect sugar and energy availability to immune signaling remain elusive.

Here, we provide insight into the regulatory mechanisms of defense responses under sugar- and energy-limited conditions. We show that SA biosynthesis and signaling are both suppressed under low sugar conditions, resulting in reduced expression of defense-related genes. In addition, SnRK1 acts as a negative regulator of defense gene expression, in part through the SA pathway, and contributes to the suppression of immunity under high humidity. Furthermore, SnRK1 physically associates with TGA transcription factors, influencing their target gene expression. These findings illuminate a central role for SnRK1 in integrating sugar and energy deprivation with immunity regulation in plants.

## Results

### Sugar limitation impairs flg22-induced defense signaling via differential effects on SA-dependent branches

Sugar availability globally influences transcriptional and metabolic reprogramming to optimize growth (32–34). To better understand the connection between sugar status and immunity, we examined PAMP responses under different sugar conditions in Arabidopsis. Wild-type (WT) seedlings grown on sugar containing medium were transferred to sugar-free liquid medium for 3 d and then treated with the bacterial flagellin-derived flg22 peptide in the presence (100 mM glucose) or absence of sugar (Fig. 1A). We first confirmed that sugar starvation markers, *DARK INDUCIBLE* 1 (*DIN1*) and *DARK INDUCIBLE* 6 (*DIN6*) (45, 66), were significantly induced under sugar-free conditions (Fig. 1B), validating sugar limitation in our experimental settings. Under sugar-sufficient conditions, the defense-related genes *NDR1/NIN1-LIKE 10* (*NHL10*) (67), *PATHOGENESIS-RELATED 1 (PR1*) and *WRKY DNA-BINDING PROTEIN 38* (*WRKY38*) were strongly induced in response to flg22 (Fig. 1B). However, their induction was reduced to less than half under sugar-free conditions, with *PR1* expression nearly abolished (Fig. 1B). Importantly, the induction of *DIN1* and *DIN6* was not affected by flg22 (Fig 1B), indicating specific downregulation of defense-inducible genes and confirming that flg22 application did not restore sugar availability. Given the SA dependence of *PR1* and *WRKY38* transcription (68, 69), we next tested whether sugar limitation affected SA-related processes. flg22 induction of *ISOCHORISMATE SYNTHASE 1* (*ICS1*), encoding a rate-limiting enzyme for defense-associated SA biosynthesis, was markedly reduced under sugar-free compared to sugar-supplemented conditions (Fig. 1B). To further address this, we examined gene expression following exogenous SA application under different sugar conditions. Intriguingly, *PR1* induction by SA treatment was abolished under sugar-free conditions, whereas *WRKY38* remained strongly induced regardless of sugar status (Fig. 1C). Together, these results suggest that the defects in flg22-induced defense responses under sugar limitation result partly from reduced SA biosynthesis, but also from impaired SA perception and/or signaling in sugar-sensitive SA-dependent branches. Thus, SA-inducible genes are not uniformly influenced by sugar availability but instead display differential sensitivity to sugar status.

**Figure 1.**
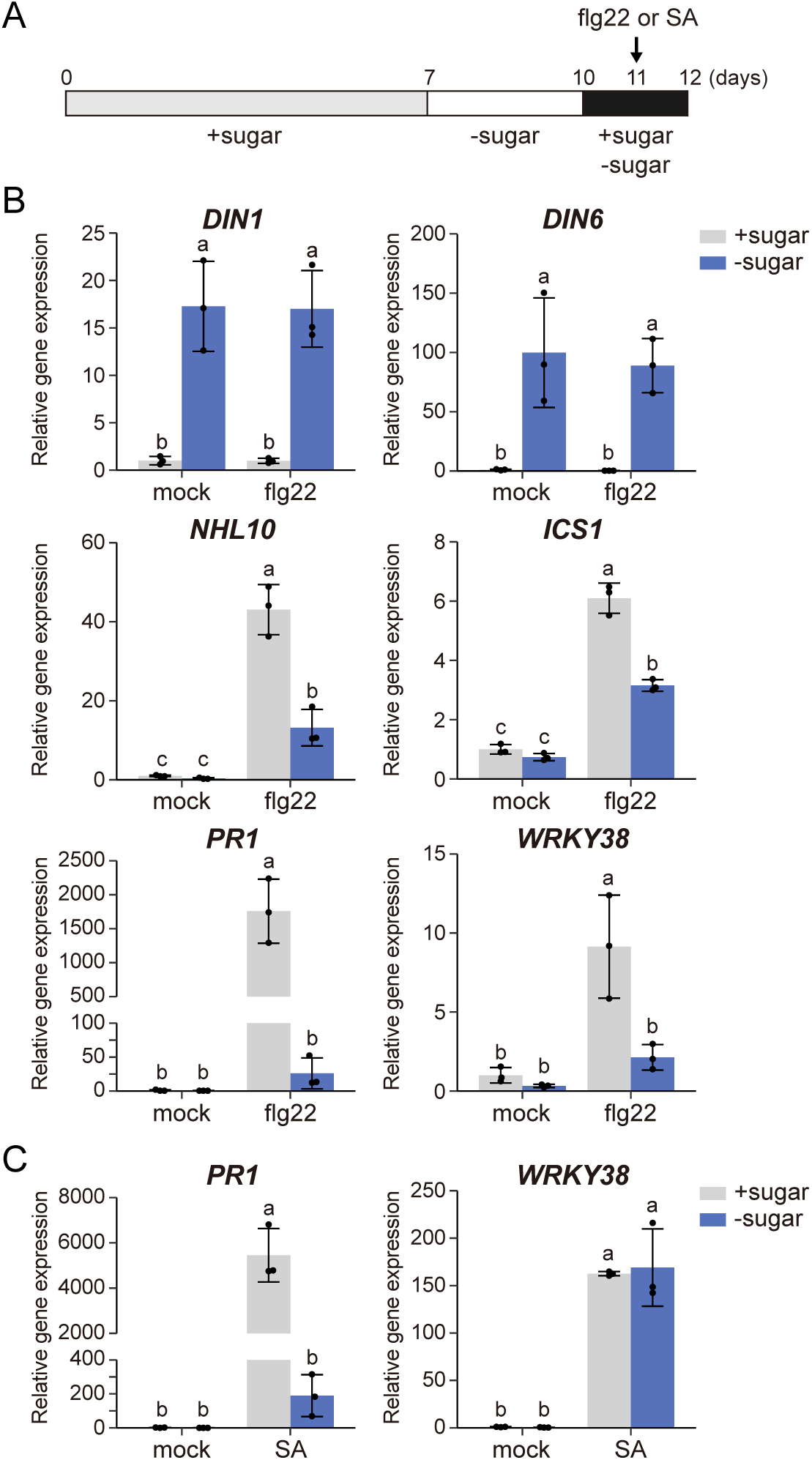
Limited sugar availability reduces the expression of defense genes. (*A*) Schematic diagram illustrating the experimental procedure for flg22 or SA treatment in the presence or absence of sugar. Seven-day-old seedlings grown on MS agar medium containing 100 mM glucose were transferred to sugar-free MS liquid medium for 3 days and then treated with glucose and elicitors. (*B*) Expression levels of sugar and energy starvation-responsive genes and defense marker genes in WT seedlings treated with mock or flg22 in the presence (+sugar) or absence (−sugar) of 100 mM glucose. (*C*) Expression levels of SA-responsive genes in WT seedlings treated with mock or SA in the presence (+sugar) or absence (−sugar) of 100 mM glucose. Expression levels were normalized to *18S rRNA* and shown relative to the mock sample with sugar. Results represent the mean ± SD (n = 3 biological replicates). Different letters indicate statistically significant differences (*P* < 0.05, Tukey’s HSD test).

### SnRK1 acts as a negative regulator of defense gene expression regardless of sugar status

As a key sensor for sugar availability, we examined whether SnRK1 is involved in the modulation of defense gene expression under sugar-limited conditions. First, we assessed SnRK1 activity under different sugar conditions in plants. While the auto-phosphorylation status of the catalytic domain T-loop has been used as a proxy for SnRK1 activity, analogous to AMPK (48, 50), its biological relevance remains unclear due to the structural differences between SnRK1 and AMPK. Therefore, we examined sugar-responsive SnRK1 activity using an ACC (Acetyl-CoA Carboxylase 1 peptide) reporter system (57), which allows for *in vivo* monitoring of SnRK1 activity in Arabidopsis. Immunoblot analysis of phosphorylated Ser79 in the ACC reporter indicated that SnRK1 activity was strongly influenced by sugar availability, significantly increasing under sugar-limited conditions (0 mM glucose) compared to sugar-supplemented conditions (100 mM glucose) (Fig. 2A).

**Figure 2.**
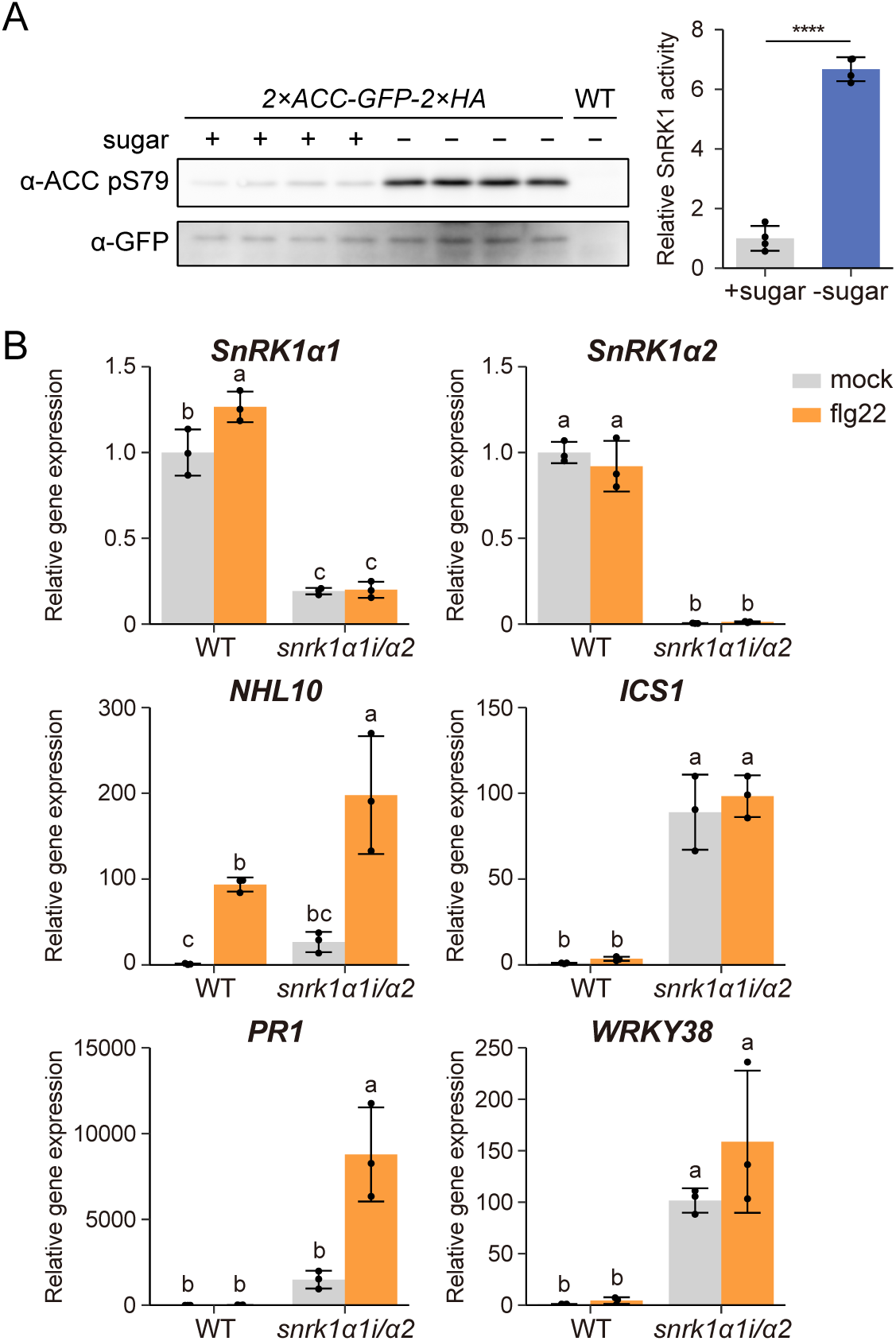
SnRK1 negatively regulates the expression of defense genes under sugar limited conditions. (*A*) *In vivo* SnRK1 activity in response to sugar availability. WT and the ACC reporter line were transferred to sugar-free medium for 3 days and then treated with 100 mM glucose (+sugar) or without glucose (−sugar) for 24 h, as described in Fig. 1A. (Left) Immunoblot analysis representing four biological replicates for each sugar condition. The immunoblot with anti-GFP antibody serves as the expression control of the ACC reporter. (Right) Quantification of SnRK1 activity using the immunoblot shown on the left. Band intensities detected with anti-ACC pS79 antibody were normalized to those with anti-GFP antibody and calculated relative to the +sugar sample. Results represent the means ± SD (n = 4 biological replicates; \*\*\*\**P* < 0.0001, Student’s *t* test). (*B*) Expression levels of defense marker genes in *snrk1α1i/α2* mutant grown under sugar limited conditions as described in Fig. 1A. WT and *snrk1α1i/α2* seedlings were pre-treated with sugar-free and DEX for 4 days, and thereafter treated with mock or flg22 for 24 h. Expression levels were normalized to *18S rRNA* and shown relative to mock-treated WT. Results represent the mean ± SD (n = 3 biological replicates). Different letters indicate statistically significant differences (*P* < 0.05, Tukey’s HSD test).

Next, we examined whether SnRK1 affects the expression of SA-dependent (*ICS1*, *PR1*, *WRKY38*) and SA-independent (*NHL10*) defense-inducible markers, using *snrk1α1i/α2* mutants, which are DEX-inducible RNAi knockdown lines for *SnRK1α1* in the *SnRK1α2-*knockout mutant (70). Under sugar-limited conditions, *WRKY38* and *ICS1* were induced even in non-elicited *snrk1α1i/α2* (Fig. 2B). Moreover, flg22 induction of *NHL10* and *PR1* was enhanced in *snrk1α1i/α2*, whereas *ICS1*, *PR1* and *WRKY38* were not significantly induced in WT (Fig. 2B). The results indicate that SnRK1 suppresses defense-related gene expression under sugar limitation.

Notably, when sugar was supplied (100 mM glucose), flg22 induction of *WRKY38* was enhanced, and *ICS1* was induced even without flg22 application in *snrk1α1i/α2* (Fig. S1). Similarly, *ICS1, PR1* and *WRKY38* were induced in non-elicited *snrk1α1i/α2* seedlings grown in a continuously sugar-supplemented medium (25 mM sucrose) (Fig. S2). We validated flg22 induction of *ICS1*, *PR1* and *WRKY38* in WT, and observed that in *snrk1α1i/α2*, three to four of the tested defense markers were induced by flg22 under both sugar supplementation regimes (Figs. S1 and S2). Notably, the marker gene expression levels were consistently higher in *snrk1α1i/α2* compared to WT when treated with flg22 (Fig. S2). These results suggest that SnRK1 functions as a negative regulator of defense gene expression and flg22-triggered immunity under both sugar-limited and sugar-sufficient conditions.

### SnRK1 broadly attenuates defense gene expression via SA/NPR1-dependent and -independent pathways

To assess an overall impact of SnRK1 on transcriptional reprogramming during pattern-triggered immunity (PTI), we performed RNA sequencing (RNA-seq) in WT and *snrk1α1i/α2* seedlings following flg22 application under sugar-limited conditions. Since prolonged sugar limitation together with reduced SnRK1 activity can cause severe growth defects and broad secondary transcriptional changes, we shortened the sugar depletion time in the RNA-seq analysis. Eight-day-old seedlings grown on sugar-sufficient medium were transferred to sugar-free medium and exposed to flg22 for 24 h (Fig. 3A and S3A). This sugar depletion regime suppressed flg22-inducible gene expression (Fig. S3B), consistent with the effects observed under the long-term depletion (Fig. 1B).

**Figure 3.**
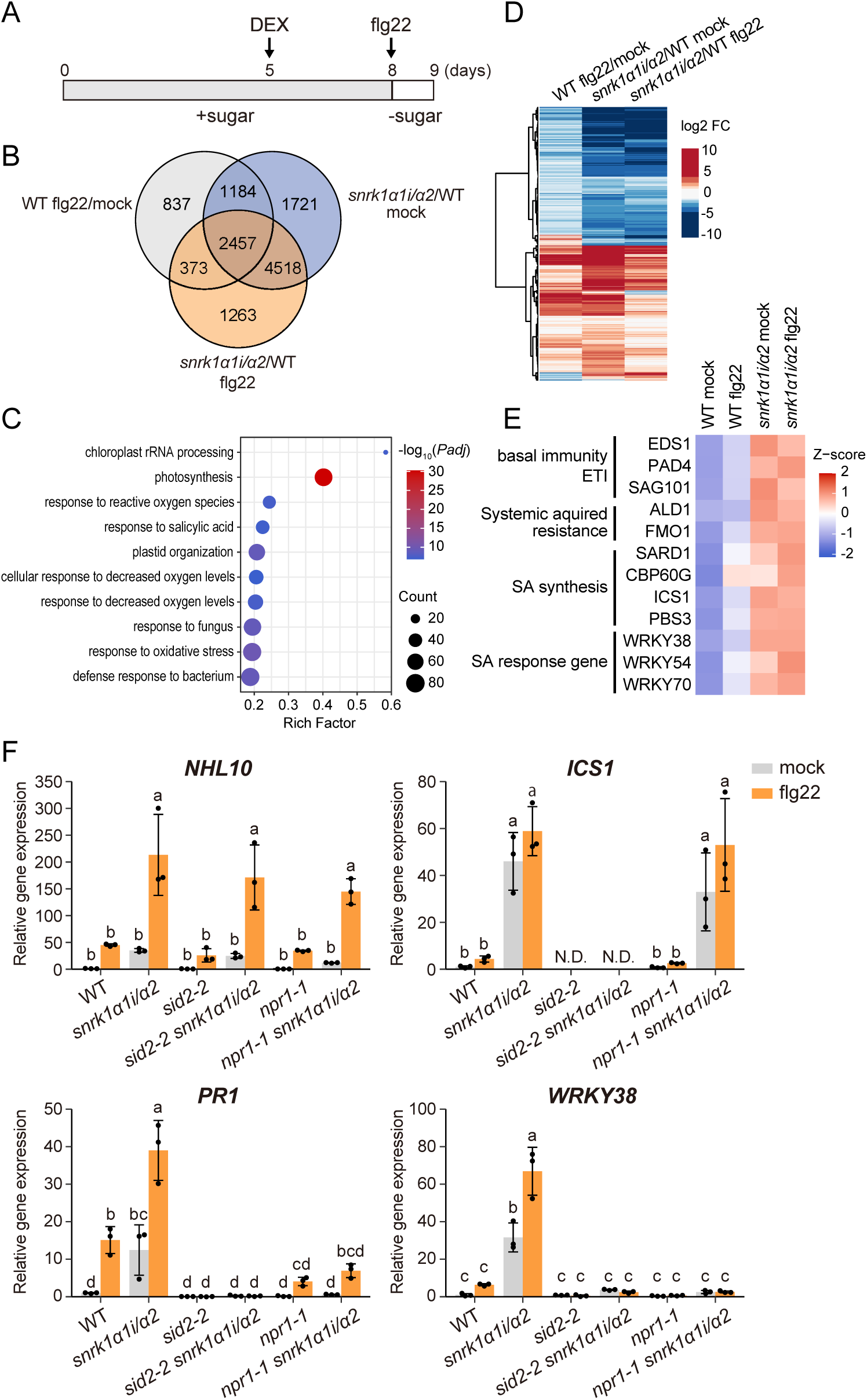
SnRK1 negatively regulates defense gene expression partially dependent on SA pathway. (*A*) Schematic diagram illustrating the experimental procedure for flg22 treatment for RNA-seq and subsequent RT-qPCR analyses. Five-day-old seedlings grown on 1/2 MS agar medium containing 25 mM sucrose were transferred to 1/2 MS liquid medium containing sucrose and DEX, and then treated with or without flg22 in the absence of sugar. (*B* to *E*) RNA-seq analysis of WT and *snrk1α1i/α2* mutant treated with mock or flg22 under sugar-limited conditions. (*B*) Venn diagram depicting the overlap of differentially expressed genes (DEGs) between WT and *snrk1α1i/α2* mutants in response to flg22. DEGs were identified with the cutoff of absolute log2 fold-change (FC) > 1 and adjusted *P* value < 0.05 (n = 4 biological replicates). (*C*) Top 10 significantly enriched GO terms of 2457 of SnRK1-regulated flg22-responsive genes identified in (*B*). (*D*) A heatmap of 2457 genes. The color key (blue to red) represents log2 FC. (*E*) A heatmap of the expression levels of representative defense-related genes in WT and *snrk1α1i/α2* mutant in response to flg22. The color key (blue to red) represents log2(TPM+1) values as z-score. (*F*) Expression levels of defense marker genes in WT, *snrk1α1i/α2*, *sid2-2*, *sid2-2 snrk1α1i/α2*, *npr1-1*, and *npr1-1 snrk1α1i/α2* mutant in response to flg22 under sugar-limited conditions, as described in (*A*). Expression levels were normalized to *18S rRNA* and shown relative to mock-treated WT. Results represent the mean ± SD (n = 3 biological replicates). Different letters indicate statistically significant differences (*P* < 0.05, Tukey’s HSD test).

Our RNA-seq analysis detected a total of 4,851 differentially expressed genes (DEGs) in WT in response to flg22, and 11,516 (1,184 + 1,721 + 2,457 + 4,518 + 373 + 1,263) DEGs in *snrk1α1i/α2* compared to WT under mock or flg22 conditions (Fig. 3B; Dataset S1). These included 3,641 (1,184 + 2,457) and 2,830 (373 + 2,457) flg22-responsive DEGs that were altered in mock and flg22-treated *snrk1α1i/α2*, respectively. Gene Ontology (GO) enrichment analysis of biological processes revealed that SnRK1-dependent flg22-responsive DEGs, especially those upregulated by flg22 and in *snrk1α1i/α2*, were enriched for immunity-related processes including SA responses, whereas those downregulated were enriched in photosynthesis-related terms (Fig. 3C and Fig. S4). Notably, 2,457 SnRK1-dependent flg22-responsive DEGs displayed similar regulation patterns both in WT in response to flg22 and in *snrk1α1i/α2* mutation under either mock or flg22 conditions (Fig. 3D). These included genes associated with SA biosynthesis and signaling, as well as key regulators of basal immunity and effector triggered immunity (ETI), such as *EDS1*, *PAD4*, and *SAG101* (71, 72), and systemic acquired resistance, such as *ALD1* and *FMO1* (73) (Fig. 3E). Collectively, these results indicate that SnRK1 constitutively acts as a negative regulator of defense gene expression, including those associated with SA.

To verify that defense enhancement in *snrk1α1i/α2* is dependent on SA, we generated triple mutants by crossing *snrk1α1i/α2* with *sid2-2* (disrupted with *ICS1* (11)) and *npr1-1* (74). Free SA levels were markedly elevated in non-elicited *snrk1α1i/α2* plants, compared to WT plants (Fig. S5), consistent with constitutive defense marker induction (Figs. 2, S1 and S2). This SA accumulation was abolished in *sid2-2 snrk1α1i/α2*, but not in *npr1-1 snrk1α1i/α2* plants (Fig. S5), indicating that SnRK1 negatively regulates ICS1-mediated SA biosynthesis. Correspondingly, the expression of *PR1* and *WRKY38* in *snrk1α1i/α2* was dramatically suppressed in both *sid2-2 snrk1α1i/α2* and *npr1-1 snrk1α1i/α2*, with or without flg22 application (Fig. 3F). By contrast, *NHL10* induction was not reduced in *sid2-2 snrk1α1i/α2* or *npr1-1 snrk1α1i/α2,* and *ICS1* expression was unaffected in *npr1-1 snrk1α1i/α2* (Fig. 3F). Together, these results indicate that SA/NPR1-dependent and SA/NPR1-independent defense pathways are both enhanced in *snrk1α1i/α2* plants.

### SnRK1 negatively regulates SA-dependent bacterial resistance

To assess the physiological relevance of our findings, we tested bacterial resistance in soil-grown *snrk1α1i*/*α2* plants. Leaves of WT and *snrk1α1i*/*α2* plants were treated with 50 μM DEX for 5 days (+DEX) and subsequently infiltrated with the biotrophic bacterial pathogen *Pseudomonas syringe* pv. *tomato* (*Pst*) DC3000 (Fig. S6A). DEX treatment efficiently suppressed *SnRK1α1* expression in *snrk1α1i*/*α2* plants without discernible phenotypic alterations (Fig. S6B). *Pst* DC3000 growth was reduced in *snrk1α1i*/*α2* plants compared with WT (Fig. 4A and B), demonstrating a negative role for SnRK1 in bacterial resistance.

**Figure 4.**
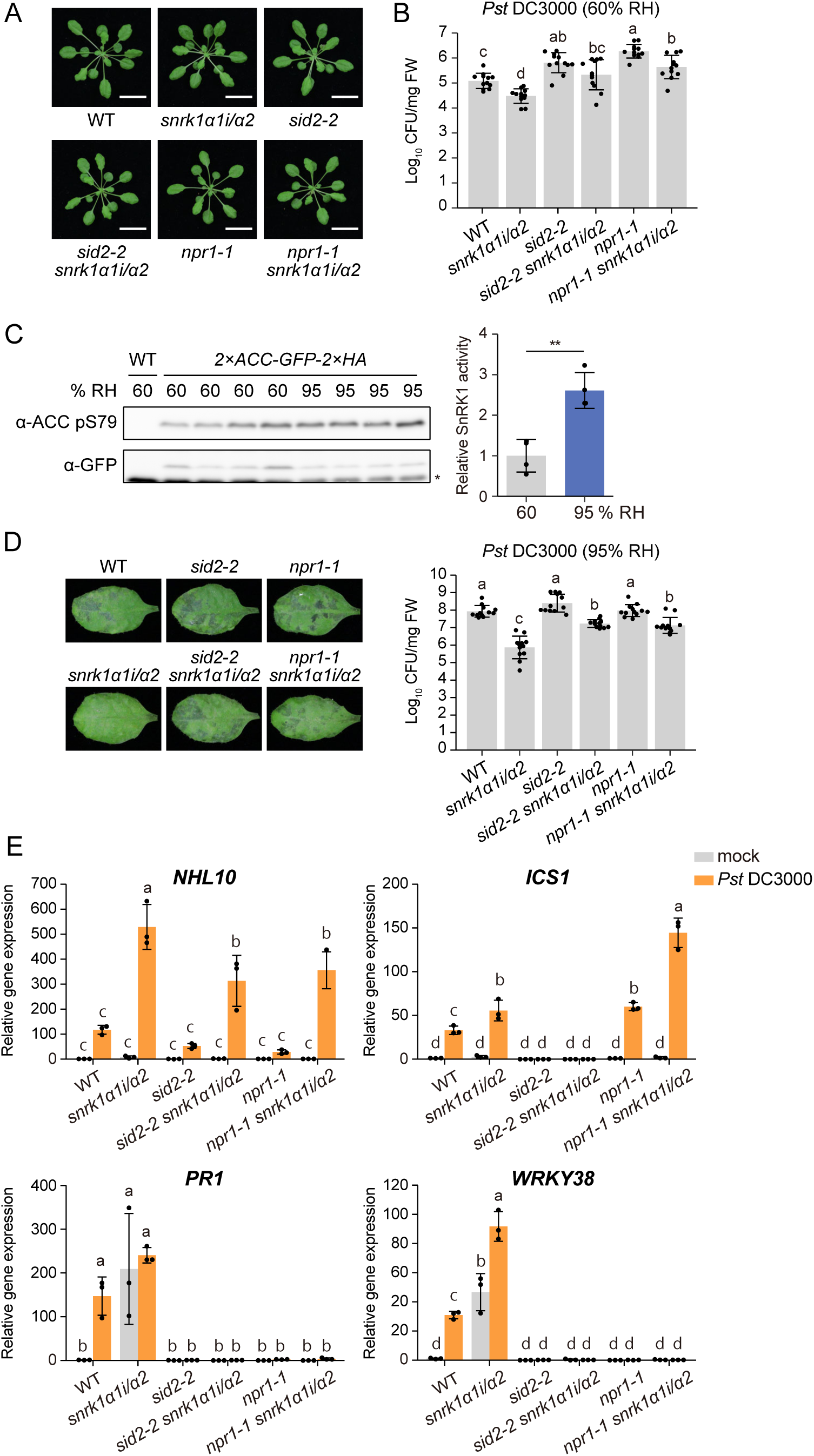
SnRK1 negatively regulates SA-mediated bacterial resistance. (A) Representative images of DEX-treated plants used in *Pst* DC3000 inoculation assays. Five-week-old soil-grown plants under 60% relative humidity (RH) were treated with DEX for 5 days. (*B*) Bacterial growth in *Pst* DC3000-inoculated leaves at 3 days post-infiltration (dpi) under moderate humidity. Plants pretreated with DEX were infiltrated with *Pst* DC3000 (OD_600_ = 0.0002) and maintained under 60% RH. Results represent the mean ± SD (n = 12 biological replicates). Different letters indicate statistically significant differences (*P* < 0.05, Tukey’s HSD). (*C*) *In vivo* SnRK1 activity under different air humidity levels. WT and the ACC reporter line were treated with moderate (60% RH) or high air humidity (95% RH) for 24 h. (Left) Immunoblot analysis representing four biological replicates for each humidity condition. The immunoblot with anti-GFP antibody serves as the expression control of the ACC reporter. An asterisk denotes nonspecific bands. (Right) Quantification of SnRK1 activity using the immunoblot shown on the left. Band intensities detected with anti-ACC pS79 antibody were normalized to those with anti-GFP antibody and calculated relative to the 60% RH sample. Results represent the means ± SD (n = 4 biological replicates). Asterisks indicate statistically significant differences (\*\**P* < 0.01, Student’s *t* test). (*D*) Disease symptoms (left) and bacterial growth (right) in *Pst* DC3000-inoculated leaves at 3 dpi under high humidity. Plants pretreated with DEX were infiltrated with *Pst* DC3000 (OD_600_ = 0.0002) and placed under 95% RH. Results represent the mean ± SD (n = 12 biological replicates). Different letters indicate statistically significant differences (*P* < 0.05, Tukey’s HSD). (*E*) Expression levels of defense-related genes in *Pst* DC3000-inoculated leaves at 1 dpi under high humidity. Plants pretreated with DEX were infiltrated with mock or *Pst* DC3000 (OD_600_ = 0.02) and placed under 95% RH. Expression levels were normalized to *ACTIN2* and shown relative to mock-infiltrated WT. Results represent the mean ± SD (n = 3 biological replicates). Different letters indicate statistically significant differences (*P* < 0.05, Tukey’s HSD test). The experimental procedure for *Pst* DC3000 inoculation and humidity treatment is described in Fig. S6A.

To determine whether this SnRK1 function relates to SA-dependent defense, we examined bacterial growth in *sid2-2 snrk1α1i*/*α2* and *npr1-1 snrk1α1i*/*α2* plants (Fig. 4A and B). Bacterial growth was increased by *sid2-2* and *npr1-1* in the *snrk1α1i/α2* background (Fig. 4B). In the *sid2-2* background, bacterial growth was no longer reduced by *snrk1α1i*/*α2*, whereas in the *npr1-1* background it remained reduced (Fig. 4B). These results indicate that SnRK1 suppresses bacterial resistance primarily by attenuating SA accumulation through ICS1, which is mediated by both NPR1-dependent and NPR1-independent pathways.

### SnRK1 contributes to defense suppression under high humidity

Plant diseases are influenced not only by host factors and pathogen virulence but also by environmental conditions (75–77). High humidity has been reported to impair SA signaling and to modulate hypoxia-related genes (28). Since SnRK1 is a central regulator of growth under energy-limited conditions including hypoxia (41–44), we examined whether SnRK1 contributes to defense suppression under high humidity. Indeed, *DIN1* and *DIN6* were upregulated under high humidity (95% relative humidity, RH) compared to moderate humidity (60% RH) (Fig. S6C), and an ACC reporter assay showed that SnRK1 activity was elevated under high humidity (Fig. 4C), together suggesting SnRK1 activation, possibly in response to cellular energy starvation, under high humidity.

We next tested whether SnRK1 regulates bacterial resistance under high humidity. WT and *snrk1α1i/α2* plants were transferred to high humidity for 3 d after *Pst* DC3000 infiltration (Fig. S6A). Bacterial growth was reduced in *snrk1α1i/α2* compared to WT even without DEX (i.e. in *snrk1α2*), and further reduced with DEX (Fig. S6D), indicating that SnRK1*α2* and SnRK1α1 act additively to suppress bacterial resistance. Bacterial growth was partially restored in *sid2-2 snrk1α1i*/*α2* and *npr1-1 snrk1α1i*/*α2* plants (Fig. 4D), indicating that SnRK1 attenuates both ICS1/SA- and NPR1-mediated resistance under high humidity. However, bacterial growth in these mutants remained lower than in *sid2-2* and *npr1-1* alone (Fig. 4D), suggesting that SnRK1 also suppresses SA/NPR1-independent resistance under high humidity.

We further examined defense marker expression during SnRK1 suppression of bacterial resistance in mature leaves. In mock-treated *snrk1α1i/α2* plants (with DEX), *PR1* and *WRKY38* were elevated compared to WT, suggesting that SA signaling was enhanced in *snrk1α1i/α2* plants also at mature stage (Fig. 4E). In addition, *WRKY38*, *NHL10* and *ICS1* were significantly up-regulated, and *PR1* also showed an increasing trend although without statistical significance, in *snrk1α1i*/*α2* compared to WT at 1 d post infiltration with *Pst* DC3000 (Fig. 4E). In *sid2-2 snrk1α1i*/*α2* and *npr1-1 snrk1α1i*/*α2*, induction of *PR1* and *WRKY38* was abolished, whereas enhanced *NHL10* induction largely remained (Fig. 4E). Together with our RNA-seq analyses (Fig. 3C-E), these results demonstrate that SnRK1 suppresses bacterial resistance by downregulating both SA/NPR1-dependent and SA/NPR1-independent defense programs.

### SnRK1 physically associates with TGAs to repress *PR1* expression

To elucidate the molecular basis of SnRK1-mediated repression of immunity under limited sugar conditions, we explored protein interactors of SnRK1. Given the sugar-dependent regulation of defense-related gene expression in response to flg22 and SA (Fig. 1B and C), we hypothesized that SnRK1 might associate with transcriptional regulators. Since *PR1* expression is directly controlled by NPR1-TGACG SEQUENCE-SPECIFIC BINDING PROTEIN (TGA) module in response to SA (15–25), we examined whether SnRK1 acts in concert with TGA transcription factors in repressing SA defense.

We first tested protein-protein interactions between SnRK1 and TGAs using a cell-free AlphaScreen assay (78, 79). TGA1-TGA7 proteins, which interact with NPR1 in Arabidopsis, are classified into three clades: clade I (TGA1, TGA4), clade II (TGA2, TGA5, TGA6) and clade III (TGA3, TGA7) (15–23). Most TGAs, except for TGA3, yielded higher AlphaScreen signals than the negative control, with particularly strong signals from clade I and clade II members (Fig. 5A and Fig. S7). Although the possible involvement of class III TGAs cannot be ruled out based solely on the AlphaScreen results, we subsequently focused on TGA4 and TGA2 as representative members of clade I and II, respectively. Consistently, yeast two-hybrid and split luciferase complementation assays in *Nicotiana benthamiana* confirmed both TGA4 and TGA2 physically interacted with SnRK1α1 (Fig. 5B and D). Moreover, yeast two-hybrid analysis revealed that the C-terminal regions of TGA4 and TGA2 are essential for their interactions with SnRK1α1 (Fig. 5C and Fig. S8).

**Figure 5.**
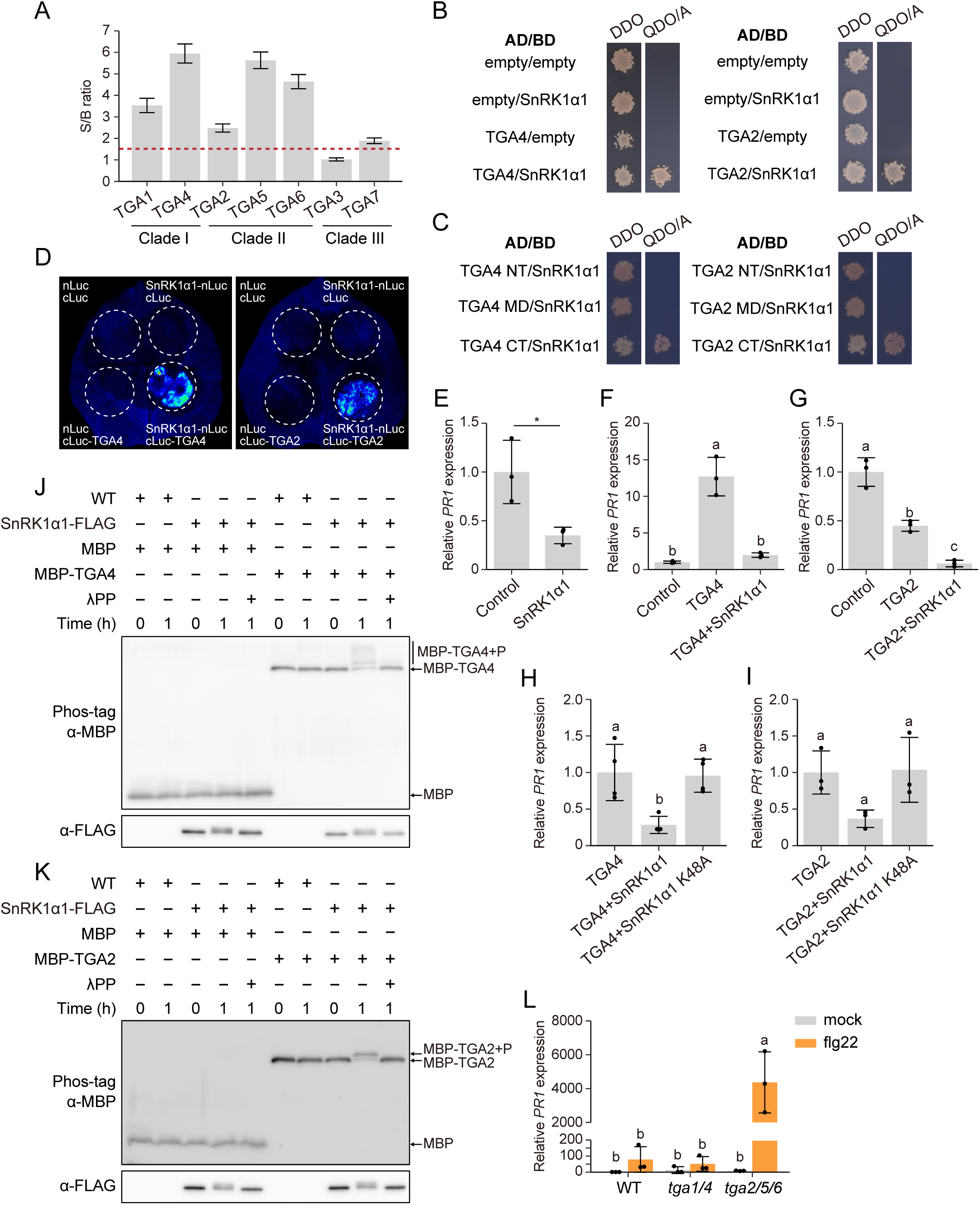
SnRK1 phosphorylates TGA transcription factors and regulates the *PR1* expression under sugar limited conditions. (A) AlphaScreen showing *in vitro* interaction of biotinylated SnRK1α1 with FLAG-TGAs. Recombinant proteins were generated in a wheat germ cell-free system. The S/B ratio was calculated by dividing the AlphaScreen signal from a wheat germ cell extract (WGE) expressing biotinylated SnRK1α1 with FLAG-TGA by the background signal, which was defined as the signal from a WGE expressing only biotinylated SnRK1α1. Results represent the mean ± SE (n = 3 biological replicates). (*B* and *C*) Yeast two-hybrid assay showing the interaction of SnRK1α1 with TGA4 and TGA2. Yeast cells were co-transformed with the indicated constructs and grown on double dropout (DDO) medium (SD-L-W) to select transformants or quadruple dropout medium (SD-L-W-H-Ade) containing 100 ng/mL Aureobasidin A (QDO/A) to test interactions. AD, transcription activation domain; BD, DNA-binding domain. The constructs used in (*C*) was described in Fig. S8A. N-terminal domain (NT), middle domain (MD), and C-terminal domain (CT). (*D*) Split luciferase complementation assay showing the interaction of SnRK1α1 with TGA4 and TGA2. Agrobacterium strains carrying the indicated constructs were mixed at a 1:1 ratio and infiltrated into *N. benthamiana* leaves. nLUC and cLUC, SnRK1α1-nLUC and cLUC, and nLUC and cLUC-TGA4 or cLUC-TGA2 were used as negative controls. (*E*) The expression level of *PR1* in Arabidopsis mesophyll protoplasts expressing *SnRK1α1*. Expression levels were normalized to *18S rRNA* and shown relative to the control. Results represent the mean ± SD (n = 3 biological replicates; \*\**P* < 0.01, Student’s *t* test). Control, protoplasts treated in the same way without plasmids. (*F* and *G*) The expression level of *PR1* in Arabidopsis mesophyll protoplasts expressing TGA4 (*F*) or TGA2 (*G*) with SnRK1α1. Expression levels were normalized to *18S rRNA* and shown relative to the control. Results represent the mean ± SD (n = 3 biological replicates). Different letters indicate statistically significant differences (*P* < 0.05, Tukey’s HSD test). (*H* and *I*) The expression level of *PR1* in Arabidopsis mesophyll protoplasts expressing TGA4 (*H*) or TGA2 (*I*) with SnRK1α1 or SnRK1α1 K48A. Expression levels were normalized to *18S rRNA* and shown relative to either the sample expressing *TGA4* (*H*) or *TGA2* (*I*). Results represent the mean ± SD (n = 4 in *H*; n = 3 in *I* biological replicates). Different letters indicate statistically significant differences (*P* < 0.05, Tukey’s HSD test). (*J* and *K*) *In vitro* phosphorylation assay. Recombinant MBP or MBP-TGA4 (*J*) or MBP-TGA2 (*K*) were incubated with immunoprecipitated proteins from WT or plants expressing *SnRK1α1-FLAG* in the presence or absence of lambda phosphatase (λPP) for the indicated time. P, phosphorylation. (*L*) The expression level of *PR1* in WT, *tga1/4*, and *tga2/5/6* mutant under sugar-limited conditions as described in Fig. 3A. Expression levels were normalized to *18S rRNA* and shown relative to mock-treated WT. Results represent the mean ± SD (n = 3 biological replicates). Different letters indicate statistically significant differences (*P* < 0.05, Tukey’s HSD test).

We next investigated the role of SnRK1 in the TGA-dependent transcriptional regulation of *PR1*. By transiently expressing *TGA4* or *TGA2* with *SnRK1α1* in mesophyll protoplast cells, we determined endogenous *PR1* expression in qPCR analyses. Consistent with the results in *snrk1α1i/α2* plants, overexpression of the *SnRK1α1* catalytic subunit suppressed *PR1* expression in this analysis (Fig. 5E and Fig. S9A). While *TGA4* overexpression activated *PR1* expression, co-expression with *SnRK1α1* significantly repressed this *PR1* activation (Fig. 5F and Fig. S9B). By contrast, *TGA2* overexpression repressed *PR1* expression, and this repression was further enhanced by *SnRK1α1* (Fig. 5G and Fig. S9C). The repression effects were diminished when a kinase-dead mutant of SnRK1α1 (SnRK1α1 K48A), substituted in the ATP-binding pocket, was co-expressed with either TGA4 or TGA2 (Fig. 5H and I and Fig. S9D and E). This indicates that SnRK1-mediated repression of *PR1* expression depends on its kinase activity.

We then examined whether SnRK1 phosphorylates TGA4 and TGA2 *in vitro*. Recombinant MBP-TGA4 and MBP-TGA2 proteins were incubated with FLAG-tagged SnRK1α1 purified from Arabidopsis plants. Phos-tag SDS-PAGE followed by immunoblot analysis revealed distinct band shifts in MBP-TGA4 and MBP-TGA2 after incubation with SnRK1 (Fig. 5J and K). These mobility shifts were eliminated by lambda phosphatase treatment (Fig. 5J and K). Furthermore, the shifts were disturbed by the AMPK kinase inhibitor Compound C (80, 81) (Fig. S10), confirming that SnRK1 phosphorylates TGA4 and TGA2. SnRK1α1 also exhibited a band shift that was sensitive to both lambda phosphatase and Compound C (Fig. 5J and K, and Fig. S10), linking SnRK1 auto-phosphorylation with its activity to phosphorylate TGA4 and TGA2.

We next evaluated the physiological relevance of the class I and class II TGAs in modulating *PR1* expression under limited sugar conditions. Class I TGAs, TGA1 and TGA4, partially redundantly promote basal resistance, with TGA1 exerting a greater effect than TGA4 (15). Class II TGAs, TGA2, TGA5, TGA6, function redundantly as bifunctional modulator of SA signaling (15, 24). Considering this genetic relationship, we examined *tga1/4* double (15) and *tga2/5/6* triple mutants (24). Interestingly, under limited sugar conditions, flg22-induced *PR1* expression was not detected in WT or *tga1/4* but was dramatically elevated in *tga2/5/6* (Fig. 5L and Fig. S11A). This upregulation was not due to reduced *SnRK1α1* and *SnRK1α2* expression in *tga2/5/6* (Fig. S11A) and was also observed under continuous sugar supplementation (Fig. S11B). Moreover, transient overexpression of *SnRK1α1* in *tga2/5/6* protoplasts failed to suppress enhanced *PR1* expression (Fig. S11C and D), suggesting that SnRK1 represses *PR1* expression at least partly through TGA2/5/6. Together, these results suggest that the class II TGAs cooperate with SnRK1 to repress *PR1* expression under both limited and sufficient sugar conditions at young seedling stage.

## Discussion

This study reveals a mechanism by which plants modulate immunity under energy and nutrient limitation through the cellular energy sensor SnRK1. We show that SnRK1 represses defense gene expression and biotrophic bacterial resistance, consistent with its established role in reprogramming transcriptional and metabolic states to optimize plant growth under energy-limited conditions, such as extended darkness, hypoxia and submergence (41–47). The increased SnRK1 kinase activity and enhanced defense gene expression in *snrk1α1i/α2* plants under sugar starvation suggest that SnRK1 actively suppresses immunity in energy- and nutrient-depleted states. Importantly, we also found the elevated defense gene expression and enhanced bacterial resistance of *snrk1α1i/α2* mutants under high humidity, indicating that SnRK1 attenuates defense responses across diverse environmental conditions.

Environmental cues are critical determinants of plant disease outcomes, together with host susceptibility and pathogen virulence, as illustrated in the “plant disease triangle” model (82). Increasing evidence highlights that abiotic stresses compromise SA biosynthesis and signaling, thereby weakening plant immunity (75–77). For instance, elevated temperatures suppress pathogen-induced SA biosynthesis and effector triggered immunity (ETI) regulators, by down-regulating the defense-related transcription factors *CALMODULIN-BINDING PROTEIN 60-LIKE G* (*CBP60g*) and *SYSTEMIC ACQUIRED RESISTANCE DEFICIENT 1* (*SARD1*) and the formation of GUANYLATE-BINDING PROTEIN-LIKE 3 (GBPL3) condensates in the nucleus (29, 83–87). Similarly, high humidity facilitates bacterial colonization by establishing water-rich apoplasts in leaves (88, 89) and reduces SA production and signaling, in part by inhibiting NPR1 ubiquitination and its binding to target gene promoters (28). Our findings suggest that SnRK1 contributes to this humidity-induced immune suppression, by inhibiting NPR1-TGA-mediated expression of defense genes, such as *PR1.* The possibility that CUL3-mediated NPR1 ubiquitination and SnRK1-mediated TGA phosphorylation converge on the NPR1-TGA module represents an intriguing direction for future research. It is also important to elucidate how SnRK1 activity is modulated in response to humidity. Given that the expression of *DIN1* and *DIN6* was increased under high humidity, changes in cellular metabolic and energy states might contribute to SnRK1 activation under high humidity. Besides, recent studies have demonstrated that a phytohormone abscisic acid (ABA) plays a crucial role in regulating plant disease responses under high humidity (89–91). Intriguingly, SnRK1 function is also regulated by ABA through the dissociation of SnRK1-SnRK2 complex (92, 93), implicating a possible involvement of ABA in modulating SnRK1 activity under high humility.

We also show that sugar limitation represses flg22/SA-induced defense gene expression in young seedlings, whereas this suppression is abolished in *snrk1α1i/α2* mutants regardless of sugar availability. Moreover, loss of *SnRK1* enhances defense gene expression and bacterial resistance in mature plants. These findings indicate that SnRK1 suppresses immunity in broad developmental and environmental contexts, potentially with its role beyond SA-dependent resistance to biotrophic pathogens.

Mechanistically, our results suggest that SnRK1 targets the NPR1–TGA transcriptional module. Class I TGAs (TGA1/4) act as activators of SA-responsive and SA biosynthesis-related genes (15, 94, 95), whereas class II TGAs (TGA2/5/6) function as dual regulators of SA-responsive gene expression. Under low SA levels (in uninfected conditions), TGA2/5/6 interact with NPR3/4 to repress defense gene expression (14). However, in pathogen-induced high-SA states, NPR3/4 dissociation promotes TGA2/5/6 interactions with NPR1, thereby activating defense gene expression (14). Consistently, our results show that TGA2/5/6 represses *PR1* expression in Arabidopsis mesophyll protoplasts under uninfected conditions. This *PR1* repression is further enhanced by SnRK1, in a manner dependent on its kinase activity. In addition, *tga2/5/6* mutants exhibited elevated flg22 induction of SA-dependent defense markers under both sugar-limited and sugar-supplemented conditions, suggesting that SnRK1 cooperates with class II TGAs to suppress immunity irrespective of sugar status. Notably, exogenous SA application is not sufficient to restore *PR1* expression under sugar starvation, implying that SnRK1-mediated repression can override SA signaling even when the latter is strengthened at young seedling stages. It should be further investigated how the sugar-starvation affects immunity at different leaf age. It is also possible that class III TGAs, TGA3 and TGA7, are associated with SnRK1 under some environmental stress.

Although the precise biochemical consequences of SnRK1-mediated TGA phosphorylation remain to be elucidated, emerging evidence suggests that post-translational regulation of TGA activity plays a critical role in integrating abiotic and biotic stress signals. For instance, INDUCER OF CBF EXPRESSION 1 (ICE1), a key transcription factor in cold stress signaling, physically interacts with TGA3 and NPR1 to promote SA-mediated defense under low temperatures (30). The brassinosteroid (BR) signaling protein kinase BR-INSENSITIVE 2 (BIN2) phosphorylates TGA4 at Ser202 or TGA3 at Ser33 to inhibit or enhance their DNA-binding activity and interactions with NPR1, respectively (96, 97). These studies highlight TGAs as molecular hubs in the transcriptional regulation of diverse stress- and growth-related signaling pathways. Our findings extend this paradigm by implicating SnRK1 as a sugar/energy-responsive regulator of TGA functions (Fig. S12).

Together, this study establishes SnRK1 as a key integrator of metabolic and environmental cues in plant immunity. By repressing SA–dependent defense and potentially other pathways, SnRK1 facilitates plant adaptation to energy deficit or humidity stress, albeit at the cost of reduced resistance. Future work should elucidate how the SnRK1-TGA module mediates sugar status-dependent defense regulation, particularly by dissecting the biochemical impact of SnRK1-mediated TGA phosphorylation on TGA-NPR1/NPR3/NPR4 interaction dynamics, and explore whether SnRK1 broadly regulates resistance to diverse pathogens.

## Acknowledgments

We thank Mie Matsubara (Nara Institute of Science and Technology) and Susumu Uehara (Nagoya University) for experimental assistance. This work was supported by Grants-in-Aid for Scientific Research from the Japan Society for the Promotion of Science (JSPS) [23H02170, 23H04184 and 25H01346 to T.S., 23K13921 to M.S., 21K14829 to S.Y., 21H02507 to Y.S., 20H05905 and 20H05906 to Y.T., respectively] and ACT-X from the Japan Science and Technology Agency (JPMJAX22BN to S.Y.). L.J. was supported by China Scholarship Council (201906350019). T.S. and M.S. were supported by Hokkaido University Young Scientist Support Program.

## Data availability

The RNA-seq data have been deposited in the DDBJ database, https://www.ddbj.nig.ac.jp (accession no. PRJDB19088).

## Author Contributions

L.J., M.S., and T.S. designed research; L.J., M.S., S.Y., K.Y., S.E., A.S., M.N., J.T. performed research and analyzed data; L.J., M.S., Y.S., and T.S. wrote the paper with inputs from S.Y. K.Y., J.T., X.F.X., and Y.T.

## Competing Interest Statement

The authors declare no conflict of interest.

---

**Fig. S1.**
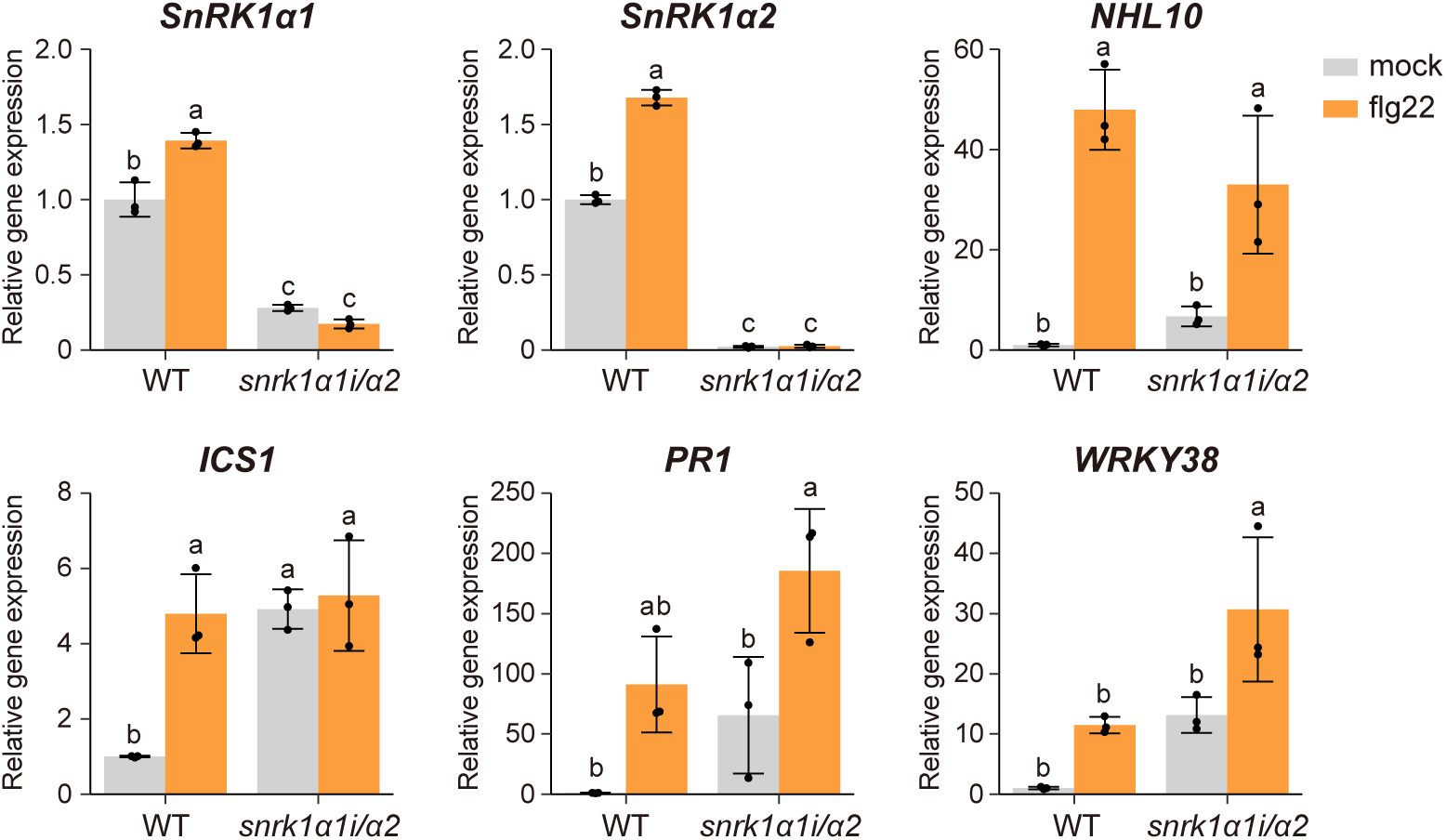
Expression levels of defense marker genes in snrk1α1i/α2 mutant under sugar-supplemented conditions. Seven-day-old WT and *snrk1α1i/α2* mutant seedlings were pretreated with sugar-free liquid medium containing DEX for 3 days, followed by transfer to medium containing 100 mM glucose and treatment with mock or flg22, as described in Fig. 1A. Expression levels were normalized to *18S rRNA* and shown relative to mock-treated WT. Results represent the mean ± SD (n = 3 biological replicates). Different letters indicate statistically significant differences (*P* < 0.05, Tukey’s HSD test).

**Fig. S2.**
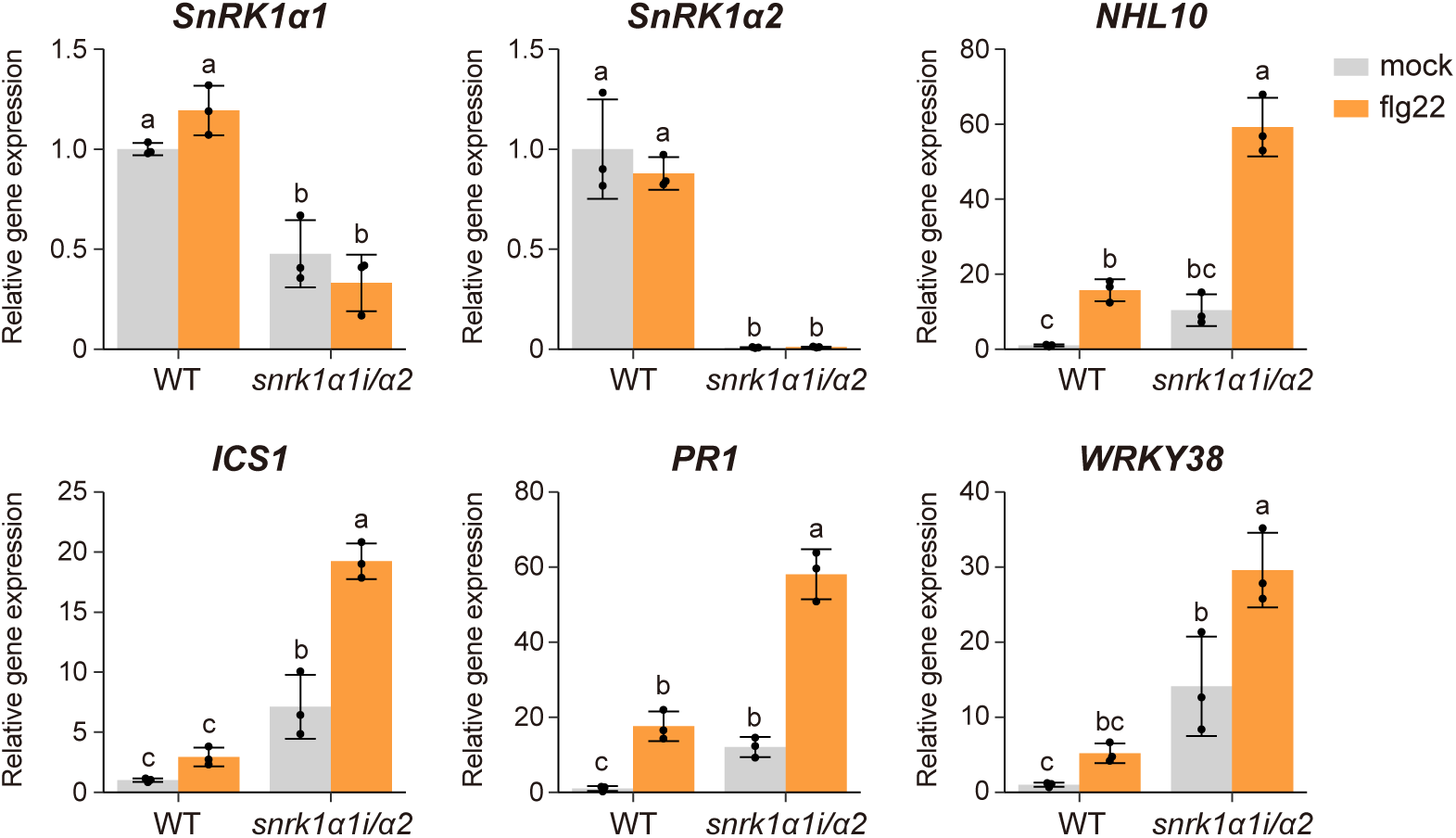
Expression levels of defense genes in *snrk1α1i/α2* mutant under continuous sugar supplementation. Five-day-old WT and *snrk1α1i/α2* mutant seedlings grown on 1/2 MS agar medium containing 25 mM sucrose were transferred to 1/2 MS liquid medium containing sucrose and DEX for 3 days, and then treated with mock or flg22 for 24 h. Expression levels were normalized to *18S rRNA* and shown relative to mock-treated WT. Results represent the mean ± SD (n = 3 biological replicates). Different letters indicate statistically significant differences (*P* < 0.05, Tukey’s HSD test).

**Fig. S3.**
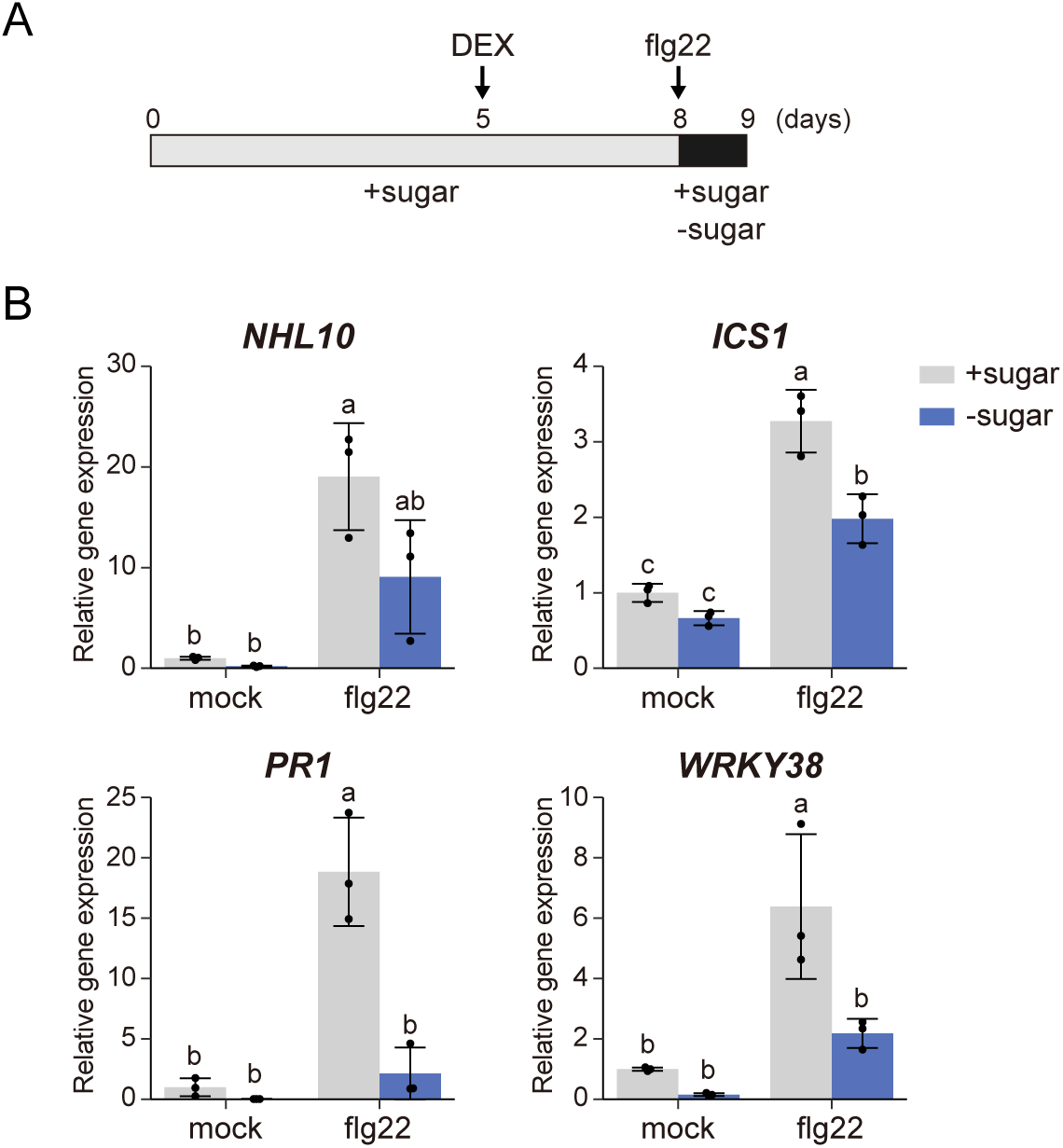
Expression levels of defense genes in WT under short-term sugar limitation. (*A*) Schematic diagram illustrating the experimental procedure for flg22 treatment in the presence or absence of sugar. Five-day-old seedlings grown in 1/2 MS agar medium containing 25 mM sucrose were transferred to 1/2 MS liquid medium containing sucrose and DEX for 3 days, and then treated with mock or flg22 for 24 h in the presence (+sugar) or absence (-sugar) of sucrose. (*B*) Expression levels of defense marker genes in WT treated simultaneously with or without sugar and flg22. WT seedlings were grown on the 1/2 MS medium containing 25 mM sucrose were treated with mock or flg22 in the presence (+sugar) or absence (−sugar) of sucrose for 24 h. Expression levels were normalized to *18S rRNA* and shown relative to the mock sample with sugar. Results represent the mean ± SD (n = 3 biological replicates). Different letters indicate statistically significant differences (*P* < 0.05, Tukey’s HSD test).

**Fig. S4.**
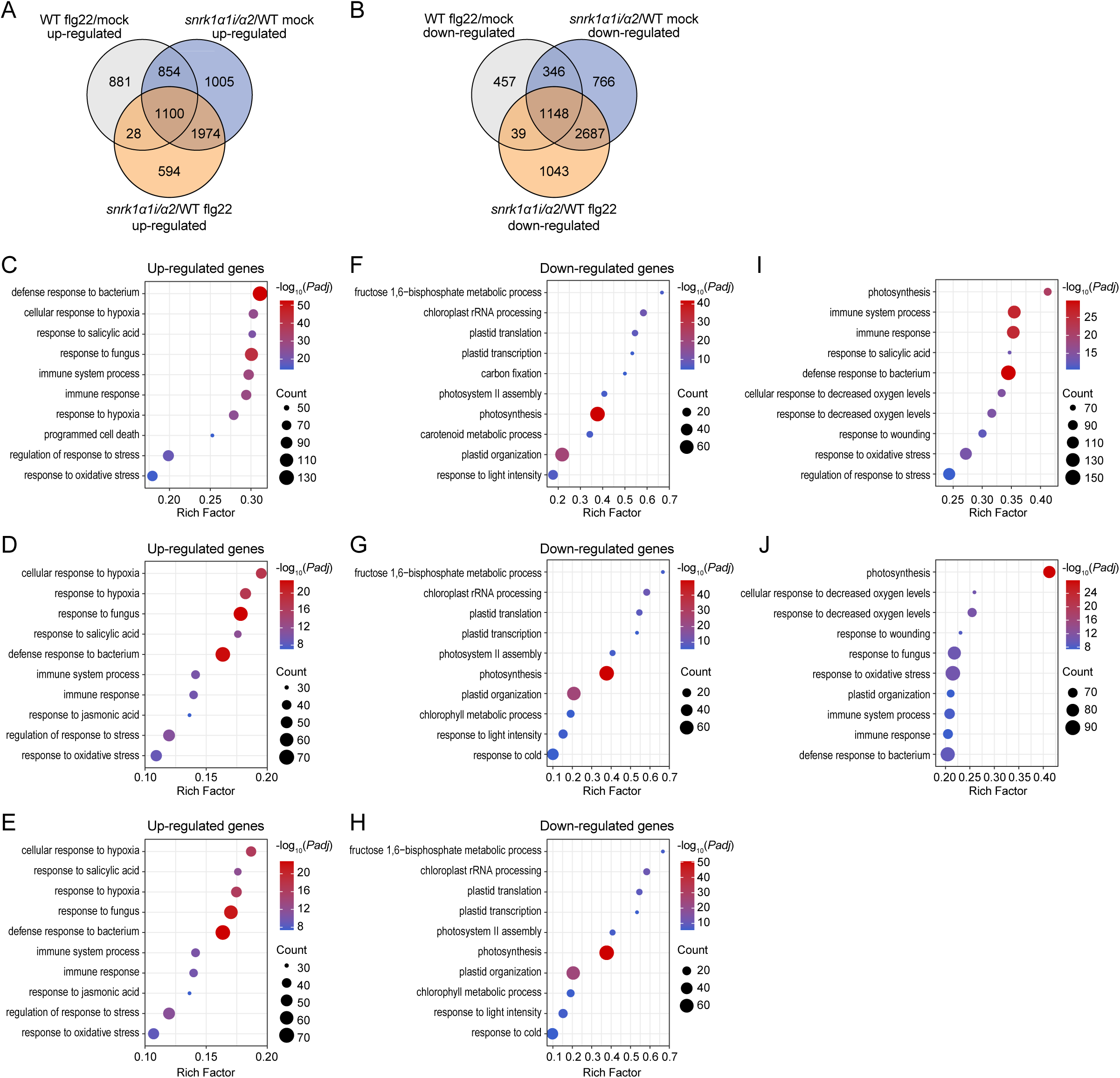
RNA-seq analysis of *snrk1α1i/α2* mutant in response to flg22. (*A* and *B*) Venn diagram depicting the overlap of up-regulated (*A*) and down-regulated genes (*B*) between WT and *snrk1α1i/α2* mutants in response to flg22. (*C* to *E*) Top 10 significantly enriched GO terms of 1954, 1128, and 1100 genes that were upregulated in the WT flg22/mock comparison and in the *snrk1α1i/α2*/WT comparison under mock (*C*), flg22 (*D*), and both conditions (*E*), as shown in (*A*). (*F* to *H*) Top 10 significantly enriched GO terms of 1954, 1128, and 1100 genes that were downregulated in the WT flg22/mock comparison and in the *snrk1α1i/α2*/WT comparison under mock (*F*), flg22 (*G*), and both conditions (*H*), as shown in (*B*). (*I* and *J*) Top 10 significantly enriched GO terms of 3641 and 2830 DEGs identified in the WT flg22/mock comparison and in the *snrk1α1i/α2*/WT comparison under mock (*I*) and flg22 (*J*), as shown in Fig. 3*B*.

**Fig. S5.**
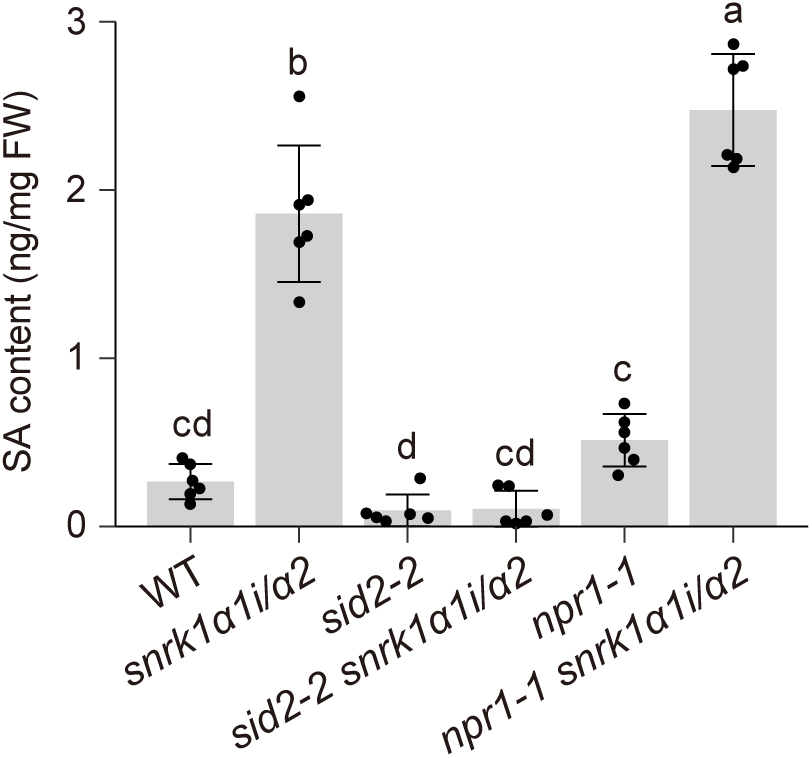
Free-SA levels in WT, snrk1α1i/α2, sid2-2, sid2-2 snrk1α1i/α2, npr1-1, and npr1-1 snrk1α1i/α2 mutant. Five-day-old seedlings grown on 1/2 MS agar medium containing 25 mM sucrose were transferred to 1/2 MS liquid media containing sucrose and DEX for 6 days. Results represent the mean ± SD (n = 6 biological replicates). Different letters indicate statistically significant differences (*P* < 0.05, Tukey’ s HSD test).

**Fig. S6.**
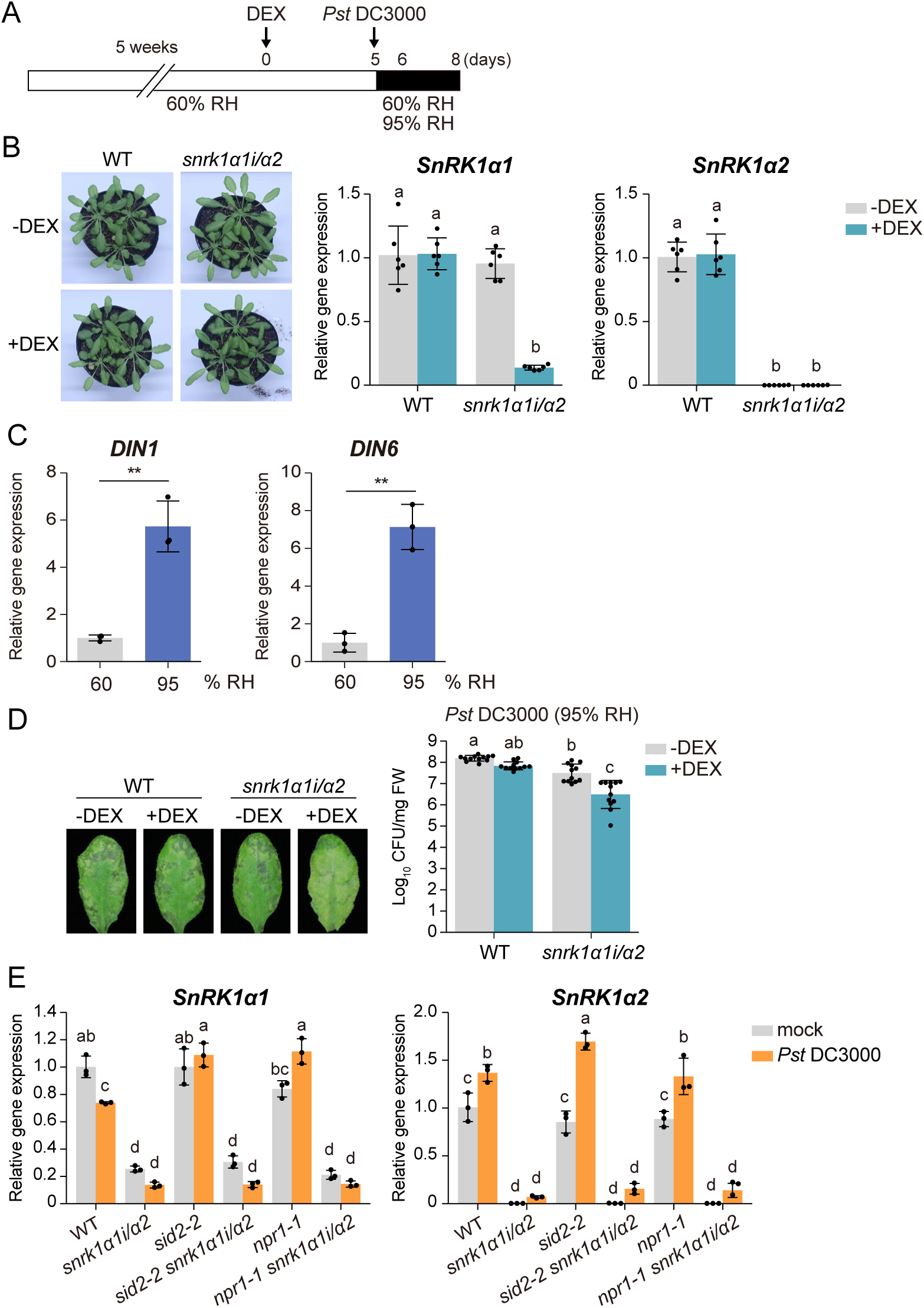
SnRK1 negatively regulates bacterial resistance. (*A*) Schematic diagram illustrating the experimental procedure for *Pst* DC3000 inoculation and humidity treatment. Five-week-old soil-grown plants under 60% relative humidity (RH) were pretreated with DEX for 5 days, followed by infiltration with *Pst* DC3000 and exposure to moderate (60% RH) or high humidity (95% RH). (*B*) Representative images (left) and expression levels of *SnRK1α* genes (right) in DEX-treated leaves. Five-week-old WT and *snrk1α1i/α2* plants under 60% RH were treated with (+) or without (−) DEX for 5 days. Expression levels were normalized to *ACTIN2* and shown relative to WT-DEX. Results represent the mean ± SD (n = 3 biological replicates). Different letters indicate statistically significant differences (*P* < 0.05, Tukey’ s HSD test). (*C*) Expression levels of sugar and energy starvation-responsive genes in the leaves of WT under different humidity levels. Five-week-old WT plants were treated with moderate (60% RH) or high air humidity (95% RH) for 24 h. Expression levels were normalized to *18S rRNA* and shown relative to 60% RH. Results represent the mean ± SD (n = 3 biological replicates). Asterisks indicate statistically significant differences (\*\**P* < 0.01, Student’ s *t* test). (*D*) Effect of DEX treatment to *Pst* DC3000 susceptibility. WT plants were pretreated with (+) or without (−) DEX, followed by inoculation with *Pst* DC3000 (OD_600_ = 0.0002) and placed under 95% RH. Disease symptoms (left) and bacterial growth (right) in *Pst* DC3000-inoculated leaves were assessed at 3 days post-inoculation (dpi). Results represent the mean ± SD (n = 12 biological replicates). Different letters indicate statistically significant differences (*P* < 0.05, Tukey’ s HSD test). (*E*) Expression levels of *SnRK1α* genes in the samples shown in Fig. 4*E*. Expression levels were normalized to *ACTIN2* and shown relative to mock-infiltrated WT. Results represent the mean ± SD (n = 3 biological replicates). Different letters indicate statistically significant differences (*P* < 0.05, Tukey’ s HSD test).

**Fig. S7.**
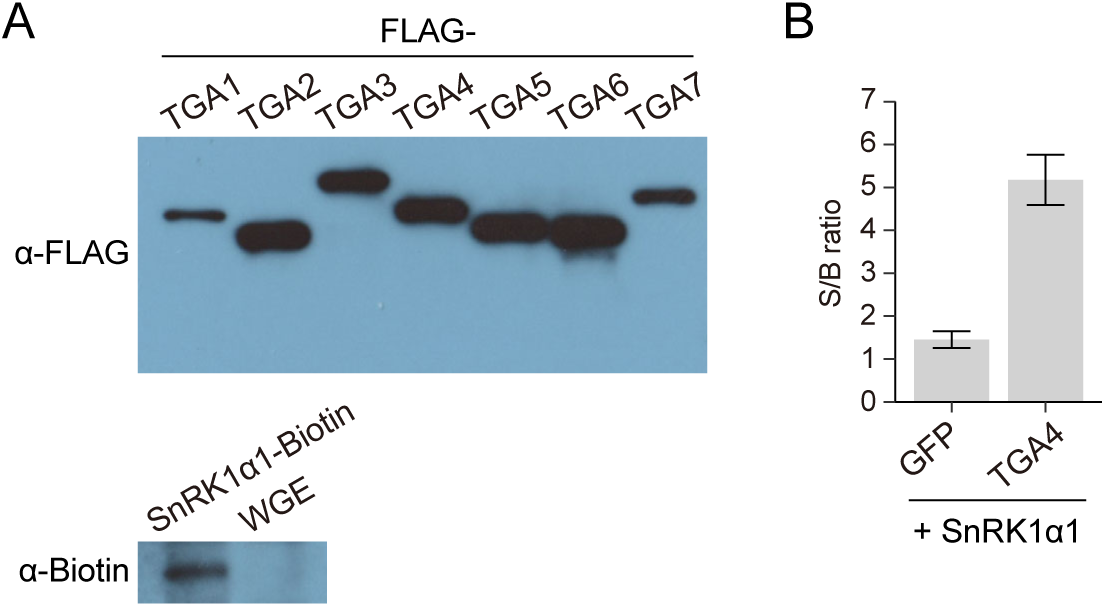
AlphaScreen assay to confirm the interaction of SnRK1α1 with TGAs. (*A*) Expression levels of recombinant proteins generated in a wheat germ cell-free system. (*B*) Biotinylated SnRK1α1 and FLAG-TGA4 generated a significant AlphaScreen signal. The S/B ratio was calculated by dividing the AlphaScreen signal from a wheat germ cell extract (WGE) expressing biotinylated SnRK1α1 with FLAG-tagged proteins by the background signal, which was defined as the signal from a WGE expressing only biotinylated SnRK1α1. FLAG-GFP was used as a negative control. Results represent the mean ± SE (n = 3 biological replicates).

**Fig. S8.**
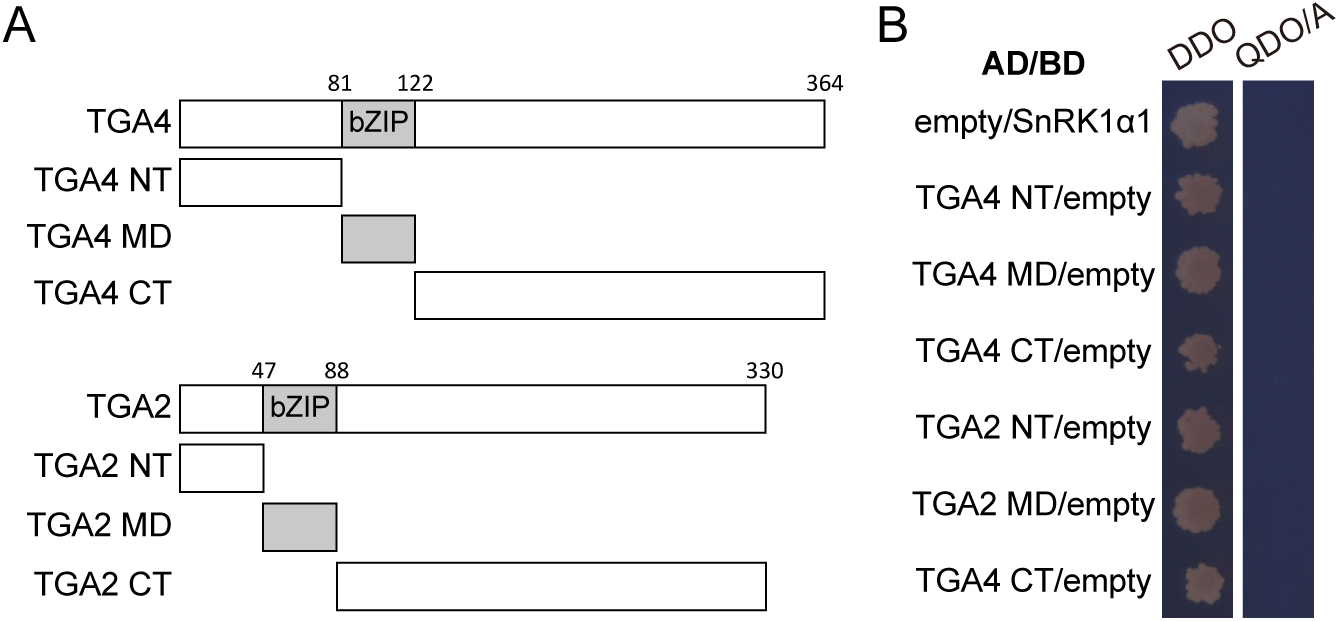
Yeast two-hybrid assay to confirm the interaction of SnRK1α1 with TGAs. (*A*) Schematic diagram of the truncated TGA used in Y2H assay shown in Fig. 5*C*. (*B*) Negative controls for Y2H assay shown in Fig. 5*C*. Yeast cells were co-transformed with the indicated constructs and grown on double dropout (DDO) medium (SD-L-W) to select transformants or quadruple dropout medium (SD-L-W-H-Ade) containing 100 ng/mL Aureobasidin A (QDO/A) to test interactions. AD, transcription activation domain; BD, DNA-binding domain. N-terminal domain (NT), middle domain (MD), and C-terminal domain (CT).

**Fig. S9.**
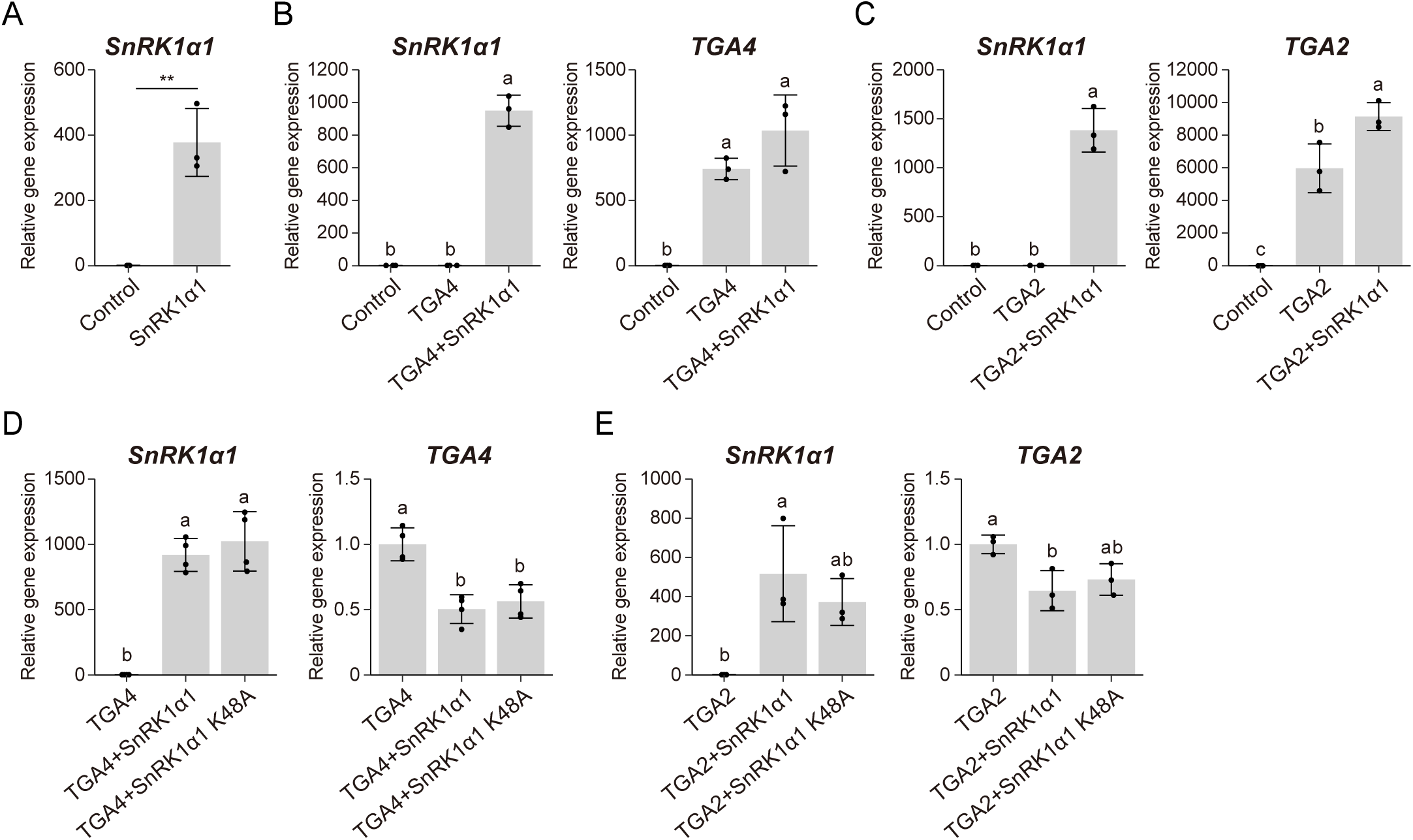
Confirmation of transient overexpression of genes in *Arabidopsis* mesophyll protoplast cells. (*A*) The expression level of *SnRK1α1* in the samples shown in Fig. 5*E*. Expression levels were normalized to *18S rRNA* and shown relative to the control. Results represent the mean ± SD (n = 3 biological replicates). Asterisks indicate statistically significant differences (\*\**P* < 0.01, Student’ s *t* test). Control, protoplasts treated in the same way without plasmids. (*B* and *C*) The expression levels of *SnRK1α1* and *TGA4*, or *TGA2* in the samples shown in Fig. 5*F* and *G*, respectively. Expression levels were normalized to *18S rRNA* and shown relative to the control. Results represent the mean ± SD (n = 3 biological replicates). (*D* and *E*) The expression levels of *SnRK1α1* and *TGA4*, or *TGA2* in the samples shown in Fig. 5*H* and *I*, respectively. Expression levels were normalized to *18S rRNA* and shown relative to either the sample expressing *TGA4* or *TGA2*. Results represent the mean ± SD (n = 4 in (*D*); n = 3 in (*E*) biological replicates). Different letters indicate statistically significant differences (*P* < 0.05, Tukey’ s HSD test).

**Fig. S10.**
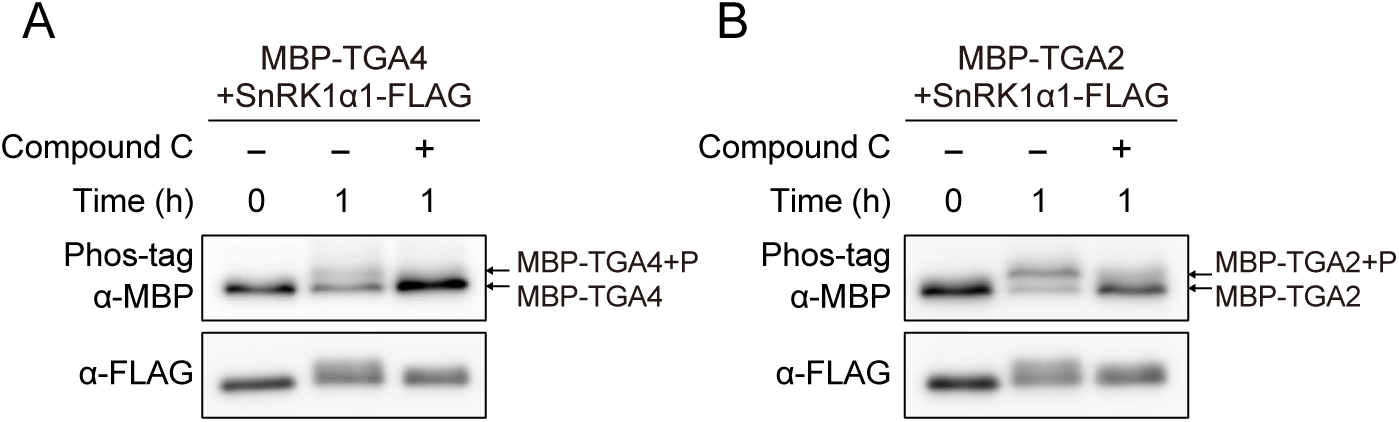
AMPK inhibitor reduced the phosphorylation of MBP-TGAs by SnRK1. (*A* and *B*) Analysis of the phosphorylation state of MBP-TGA4 (*A*) or MBP-TGA2 (*B*) incubated with immunoprecipitated proteins from plants expressing SnRK1α1-FLAG in the presence (+) or absence (−) of the AMPK inhibitor compound C for the indicated time. P, phosphorylation.

**Fig. S11.**
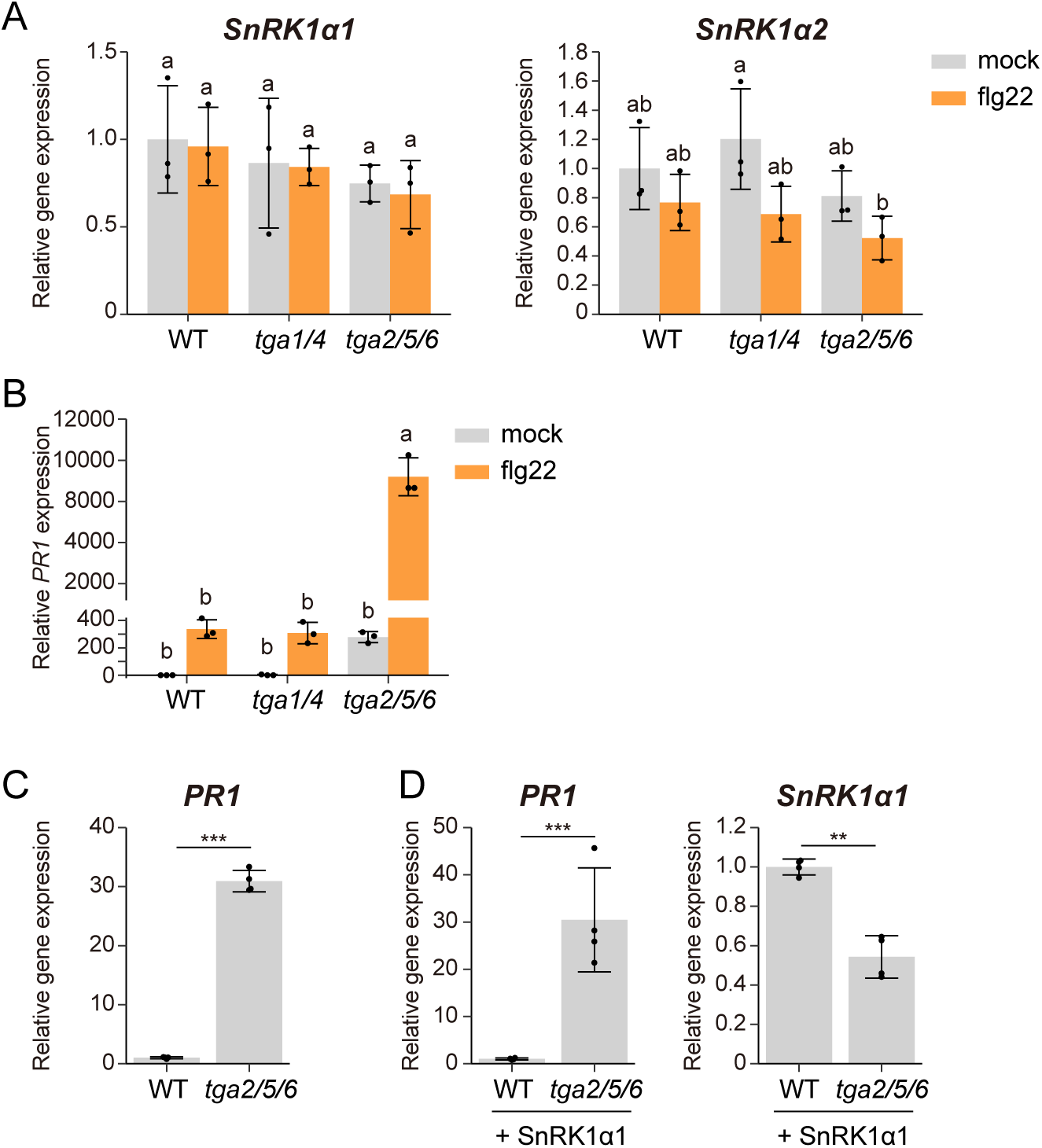
Defense-related gene expression in *tga* mutants. (*A*) The expression level of *SnRK1α* genes in the samples shown in Fig. 5*L*. Expression levels were normalized to *18S rRNA* and shown relative to mock-treated WT. Results represent the mean ± SD (n = 3 biological replicates). Different letters indicate statistically significant differences (*P* < 0.05, Tukey’ s HSD test). (*B*) The expression level of *PR1* in WT, *tga1/4*, and *tga2/5/6* mutant grown under continuous sucrose supplementation. Five-day-old seedlings grown on 1/2 MS agar medium containing 25 mM sucrose were transferred to 1/2 MS liquid medium containing sucrose for 3 days, and then treated with mock or flg22 for 24 h. Expression levels were normalized to *18S rRNA* and shown relative to mock-treated WT. Results represent the mean ± SD (n = 3 biological replicates). Different letters indicate statistically significant differences (*P* < 0.05, Tukey’ s HSD test). (*C* and *D*) The expression level of *PR1* in *Arabidopsis* mesophyll protoplasts isolated from WT or *tga256* mutant and transfected with no vector (*C*) or SnRK1α1 (*D*). Expression levels were normalized to *18S rRNA* and shown relative to the WT sample. Results represent the mean ± SD (n = 4 biological replicates; \*\*\**P* < 0.001, \*\**P* < 0.01, Student’ s *t* test).

**Fig. S12.**
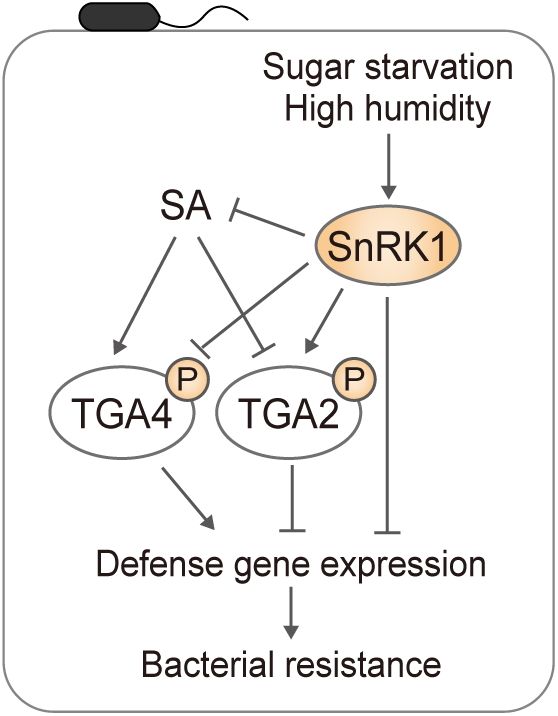
A hypothetical model illustrating the modulation of bacterial resistance by SnRK1 under energy-limited conditions. Under energy-limited conditions, such as sugar starvation and high humidity, activation of SnRK1 leads to reduction in bacterial resistance through SA-dependent and SA-independent pathways. SnRK1 negatively regulates ICS1-mediated SA biosynthesis and represses defense gene expressions by phosphorylating TGA transcription factors.

## SI Materials and Methods

### Plant materials and growth conditions

*Arabidopsis thaliana* Columbia-0 (Col-0) ecotype was used as wild type (WT) in this study. The ACC reporter line and the transgenic plants expressing *SnRK1α1-FLAG* (1), *snrk1α1i/α2* (2), *sid2-2* (3), *npr1-1* (4), *tga1/4* (*tga1-1 tga4-1*) (5), and *tga2/5/6* (6) were previously described. The *sid2-2 snrk1α1i/α2* and *npr1-1 snrk1α1i/α2* plants were obtained by crossing either *sid2-2* or *npr1-1* with *snrk1α1i/α2*.

All seeds were surface-sterilized and stratified at 4°C in the dark for 2 days. Plants were grown at 22°C with white fluorescent light provided by full-spectrum white fluorescent light bulbs (FL40SW; NEC) with a fluence rate of 60–80 μmol m^-2^ s^-1^. Unless otherwise specified, plants were grown on 1/2 Murashige and Skoog (MS) agar medium containing 25 mM sucrose and 0.5 g/L MES (pH 5.7).

To analyze the effects of sugar on gene expressions and SnRK1 activity after relatively long-term sugar limitation, plants were grown under continuous light on MS agar medium containing 100 mM glucose for 7 days and then were transferred to sugar-free MS liquid medium for 3 days with agitation at 100 rpm prior to sugar treatment. For elicitor treatment, plants were thereafter treated with 1 μM flg22 (dissolved in sterile water) or 10 mg/L SA (Sigma-Aldrich; dissolved in 0.1% dimethyl sulfoxide (DMSO)) for 24 h in the presence or absence of 100 mM glucose (see Fig. 1*A*).

For RNA-seq and subsequent RT-qPCR analyses using *snrk1α1i/α2*, the sugar-free treatment period was shortened to avoid growth defects and their secondary effects on global gene expression. Plants were grown under long-day conditions (16 h light/8 h dark) on 1/2 MS agar medium containing 25 mM sucrose for 5 days and then transferred to ½ MS liquid medium containing 25 mM sucrose supplemented with 10 μM dexamethasone (DEX) (dissolved in 0.1% EtOH) for 3 days with agitation at 100 rpm. Subsequently, plants were treated with 1/2 MS liquid medium containing 1 μM flg22 for 24 h in the presence or absence of sucrose (see Fig. 3*A*, *SI Appendix,* Fig. S3*A*).

For *Pst* DC3000 inoculation assay, plants were grown in a soli mixture [50% Super Mix A (Sakata Seed), 50% Vermiculite G20 (Nittai)] under short-day conditions (22℃, 8 h light/16 h dark, 60% relative humidity (RH)) for 5 weeks. Subsequently, leaves were infiltrated with 50 μM DEX (dissolved in 0.5% EtOH) for 5 days prior to *Pst* DC3000 inoculation.

### *Pst* DC3000 inoculation assay

*Pst* DC3000 was cultured overnight in King’s B liquid medium [2% Bacto Proteose Peptone No. 3 (Gibco, cat no. 211693), 0.15% K_2_HPO_4_, 1% glycerol, 5 mM MgSO_4_] containing 50 μg/mL rifampicin at 28°C with shaking at 180 rpm. Bacteria were collected by centrifugation at 2,000 g for 10 min at 25°C, washed once with sterile water, and resuspended in sterile water to OD_600_ = 0.0002 and 0.2 for bacterial growth and gene expression analyses, respectively. The bacterial suspensions were infiltrated into the DEX-treated leaves (3 leaves per plant), and excess solution was removed with tissue paper. The infiltrated plants were kept at 60% RH for 1 h to allow evaporation of the infiltrated solutions, and then maintained at 60% RH (moderate humidity) or 95% RH (high humidity) by covering with a clear plastic container. For bacterial growth quantification, three inoculated leaves were collected from three different plants at 3 days post-inoculation (dpi) and ground in 1 mL of sterile water. 10 μL aliquots of serial dilutions (10^-1^ to 10^-8^) were spotted onto NYGA medium [0.5% Bacto Peptone (Gibco, cat no. 211677), 0.3% yeast extract (Nacalai Tesque, cat no. 15838-45), 2% glycerol, 1.5% agar (Nacalai Tesque, cat no. 01162-15)] containing 50 μg/mL rifampicin. Colony forming units (CFU) were counted 2 days of incubation at 28°C. For gene expression analysis, three inoculated leaves were collected from three different plants at 1 dpi.

### Free SA quantification

Plants were grown under long-day conditions on 1/2 MS agar medium containing 25 mM sucrose for 5 days and then were transferred to 1/2 MS liquid medium containing 25 mM sucrose supplemented with 10 μM DEX for 6 days. Extraction and quantification of free SA by gas chromatography-mass spectrometry as described previously (7).

### Transient expression in leaf mesophyll protoplasts

To construct vectors for protoplast transformation, pENTR/D-TOPO constructs harboring the TGA2 or TGA4 CDS were recombined into pUGW2 (8) using Gateway LR clonase II (Invitrogen). For expressing SnRK1α1, the pUGW2 vector harboring either the wild-type SnRK1α1 or the SnRK1α1 K48A CDS (1) was used. To isolate leaf mesophyll protoplasts, plants were grown on the MS agar medium containing 1% sucrose for 2 weeks and then were transferred to soil for 4 weeks under short-day conditions (8 h light/16 h dark). Plasmids were introduced by polyethylene glycol mediated transformation. After transfection, protoplasts were incubated at 22℃ for 14-16 h under dim white light before harvesting.

### Transcript analysis

Total RNA was extracted using TRIzol reagent (Invitrogen) or Sepasol-RNA I Super G (Nacalai Tesque), and treated with RQ1 RNase-free DNase (Promega) according to the manufacturer’s protocols. First-strand cDNA was synthesized using oligo(dT) primer (Promega) and ReverTraAce reverse transcriptase (Toyobo) or PrimeScrip RT reagent Kit with gDNA Eraser (TaKaRa). Quantitative real-time PCR (qRT-PCR) was performed on either AriaMx real-time PCR system (Agilent Technologies) with TB Green Premix Ex Taq II (TaKaRa) or Thermal Cycler Dice Real Time System III (TaKaRa) with Power SYBR Green PCR Master Mix (Applied Biosystems). Gene-specific primers used for qRT-PCR are listed in *SI Appendix,* Table S1.

### RNA-seq and data analysis

600 ng of total RNA extracted using RNeasy Plant Mini Kit (Qiagen) was used for library preparation. PolyA selected and strand-specific library was prepared using KAPA mRNA HyperPrep Kit (cat. no. KK8580, Kapa Biosystems) with KAPA Universal adaptor (cat. no. 9063781001, Kapa Biosystems) and KAPA Unique Dual Index Primer Mixes (cat. no. 9134336001, Kapa Biosystems) following the manufacturer’s protocol. Four biological replicates of each genotype under each growth condition were subjected to sequencing. A total of sixteen libraries were pooled and 150 bp paired-end sequenced by the HiSeqX Ten sequencer (Illumina). Quality control and pre-processing of raw paired-end reads were performed using fastp v0.19.5, an ultra-fast FASTQ preprocessor (9). Reads were aligned to the Arabidopsis TAIR10 reference genome using STAR v2.7.10b (10). The number of reads mapped to each gene was counted using featureCounts v2.0.1. (11). Differential expression analysis was performed using DESeq2 in R (12). Genes with false discovery rate (FDR) adjusted *p*-value < 0.05 with the cutoff set to |log_2_ fold-change (FC)| > 1 in each comparison were identified as differentially expressed genes (DEGs). The GO enrichment analysis of DEGs was performed using clusterProfiler v4.6.0 (13) in R. The transcripts per million (TPM) value of each gene was calculated using TPMCalculator v.0.0.3 (14).

### *in vivo* SnRK1 activity assay

Proteins were extracted in 1×SDS sample buffer and were denatured for 5 min at 95°C. The supernatants obtained by centrifugation at 20,000 g for 5 min at 25°C were subjected to immunoblot analysis. Relative SnRK1 activity was determined by the quantification of immunoblot signals detected by anti-ACC pS79 antibody normalized to those detected by anti-GFP as a reporter expression control.

### Recombinant protein expression and purification

To construct the vectors for recombinant protein expression, pENTR/D-TOPO constructs harboring the TGA2 or TGA4 CDS were recombined into pDEST-mal (15). The constructs were expressed in the *Escherichia coli* strain BL21(DE3) pLysS. Bacteria from a saturated culture were inoculated into fresh LB media at 37°C with agitation at 220 rpm until the OD_600_ reached approximately 0.6. Isopropyl β-D-thiogalactopyranoside and ethanol were added to the suspensions to final concentrations of 0.2 mM and 2%, respectively. After incubation for 16 h at 18°C with agitation at 160 rpm, cells were collected and resuspended in lysis buffer [50 mM Tris-HCl (pH7.5), 100 mM NaCl, 10% Glycerol] supplemented with Complete Protease Inhibitor Cocktail (Roche), 1 mM EDTA and 1 mM DTT. The cell lysates were clarified by ultracentrifugation at 80,000 g for 1 h at 4°C. MBP-fused proteins were purified with amylose resin (New England Biolabs) according to the manufacturer’s instructions.

### *in vitro* phosphorylation assay

For the preparation of SnRK1α1-FLAG proteins, WT and *SnRK1α1-FLAG* plants were grown on 1/2 MS agar medium containing 25 mM sucrose for 16 days, and then the shoots were harvested. The immunoprecipitation was performed as described previously (1). Briefly, frozen and ground samples were homogenized with the extraction buffer [50 mM Tris-HCl (pH7.5), 150 mM NaCl, 10% glycerol, 10 mM MgCl_2_, 1 mM EDTA, 0.5% Triton-X-100, 10 μM MG132, Complete Protease Inhibitor Cocktail (Roche), and PhosSTOP (Roche)]. Crude extracts were subjected to immunoprecipitation with anti-FLAG M2 magnetic beads (Millipore, cat no. M8823) for 1 h at 4℃. The bound proteins were eluted with elution buffer [150 μg/ml 3×FLAG peptides (Sigma-Aldrich, cat.no. F4799), 50 mM Tris-HCl (pH7.5), 10 mM MgCl_2_, 0.1% Triton-X-100] for 1 h at 4℃.

For the *in vitro* phosphorylation assay, SnRK1α1-FLAG purified from the plants and 500 ng of MBP-TGAs or MBP were incubated in kinase reaction buffer [50 mM Tris-HCl (pH7.5), 10 mM MgCl_2_, 2 mM DTT, 1 mM ATP, 10 μM MG132, Complete Protease Inhibitor Cocktail (Roche), and PhosSTOP (Roche)] for 1 h at 30℃. Reactions were stopped by adding 2×SDS sample buffer, and the samples were denatured for 5 min at 95℃. For assays using the AMPK kinase inhibitor, SnRK1α1-FLAG and 500 ng of MBP-TGAs were incubated in the kinase reaction buffer supplemented with 50 μM Compound C (Selleck Chemicals, cat. no. S7840; dissolved in 1% DMSO) and 200 μM ATP. For the *in vitro* dephosphorylation assay, SnRK1α1-FLAG and 500 ng of MBP-TGAs or MBP were incubated in the kinase reaction buffer supplemented with 200 units of lambda phosphatase (New England Biolabs, cat. no. P0753) for 1 h at 30℃.

### Phos-tag SDS-PAGE

Proteins were separated by Mg^2+^ based Phos-tag SDS-PAGE, following the manufacturer’s protocol (Wako). The 7.5% polyacrylamide gel containing 50 mM Phos-tag Acrylamide AAL-107 (Wako) was used. The gel was washed twice in Tris-Glycine transfer buffer containing 10 mM EDTA for 10 min before blotting.

### Immunoblot analysis

Proteins were separated by SDS-PAGE and transferred to a PVDF membrane. The membranes were blocked with 5% non-fat milk and were incubated with the following antibodies. Anti-ACCpS79 (ACC phosphor-Ser79 specific monoclonal antibody D7D11) (1:1000, MBL, cat. no. 11818), anti-GFP (1:1000, MBL, cat. no. 598), anti-MBP (1:5000, New England Biolabs, cat. no. E8032), and anti-FLAG (1:1000, MBL, cat. no. PM020) were used as a primary antibody. Anti-Mouse IgG (HRP-Linked Whole Ab Sheep, GE Healthcare, cat. no. NA931V) and anti-Rabbit IgG (Cell Signaling Technology, cat. no. 7074) were used as a secondary antibody. To improve immunodetection, antibodies were diluted in Can Get Signal Immunoreaction Enhancer Solution (TOYOBO).

### AlphaScreen analysis

To explore the interaction between SnRK1α1 and TGAs, proteins were synthesized with N-terminal FLAG-tagged TGAs and N-terminal biotinylated SnRK1α1 (Biotin-SnRK1α1) using the Cell-Free Protein Synthesis Kit (BioSieg) as previously described (16). The biotinylated proteins were dialyzed against PBS buffer [137 mM NaCl, 8.1 mM Na_2_HPO_4_, 2.68 mM KCl, 1.47 mM KH_2_PO_4_] at 4°C for 24 h. The synthesized proteins were confirmed by immunoblotting with an anti-FLAG antibody (Wako, cat. no. 014-22383) and streptavidin-HRP (Cell Signaling Technology, cat. no. 3999). To evaluate the interactions between FLAG-tagged TGAs and Biotin-SnRK1α1, the AlphaScreen™ system [FLAG (M2) Detection Kit, PerkinElmer] was performed as described previously (17). Briefly, 25 µl of reaction buffer [1xControl buffer, 0.01% Tween 20 (Sigma Aldrich), 0.1% BSA (Sigma Aldrich), 500 ng anti-FLAG acceptor beads, 500 ng streptavidin-coated donor beads, produced FLAG-tagged protein and biotinylated protein] was incubated in a 384-well plate at 22°C for 12 h. AlphaScreen units were measured using an EnSight® Multimode Plate Reader (PerkinElmer).

### Yeast two hybrid

To construct the vectors for Y2H assays, pENTR/D-TOPO constructs harboring the TGA2 CDS, TGA4 CDS, their truncated forms, or the SnRK1α1 CDS (1) were recombined into pGADT7-GW (a gift from Yuhai Cui; Addgene plasmid # 61702; http://n2t.net/addgene:61702; RRID:Addgene_61702) or pGBKT7-GW (a gift from Yuhai Cui; Addgene plasmid # 61703; http://n2t.net/addgene:61703; RRID:Addgene_61703) (18), respectively. To test interactions between TGAs and SnRK1α1, pGADT7 and pGBKT7 derived constructs were co-transformed into the Y2H Gold strain of yeast (Clontech, cat no. 630498) using a Frozen-EZ Yeast Transformation II Kit (Zymo Research) according to the manufacturer’s protocol. Yeast cells were grown for 3 days at 28 °C on double dropout (DDO) media [SD/-Tryptophan (W)/-Leucine (L)] to select transformants and on quadruple dropout (QDO/A) medium [SD/-Tryptophan (W)/-Leucine (L)/-Histidine (H)/-Adenine (Ade)] containing 100 ng/mL Aureobasidin A (AbA) to test interactions. The combination with the complementary empty vectors pGADT7 (Clontech, cat no. 630442) and pGBKT7 (Clontech, cat no. 630443) were used as a negative control.

### Split-luciferase complementation assays

To construct vectors for split-luciferase complementation assays, the TGA2 or TGA4 CDS were cloned into pJW772 (19) with a C-terminal LUC (cLUC), and the SnRK1α1 CDS was cloned into pJW771 (19) with an N-terminal LUC (nLUC). *Agrobacterium tumefaciens* strain GV3101 harboring the plasmids was cultured overnight at 30°C in liquid LB medium. Bacteria were collected and resuspended in infiltration buffer [10 mM MgCl_2,_ 10 mM MES-NaOH (pH5.7), 200 μM acetosyringone] to OD_600_ = 0.2. After incubation at room temperature for 2 h with gentle agitation, the suspensions were infiltrated into *Nicotiana benthamiana* leaves. At 48 h post infiltration, leaves were infiltrated with luciferase substrate solution (Promega) and kept in dark for 6 min. Luminescence images were captured using a high resolution, low illumination digital cold camera (Tanon, China).

**Table S1.**
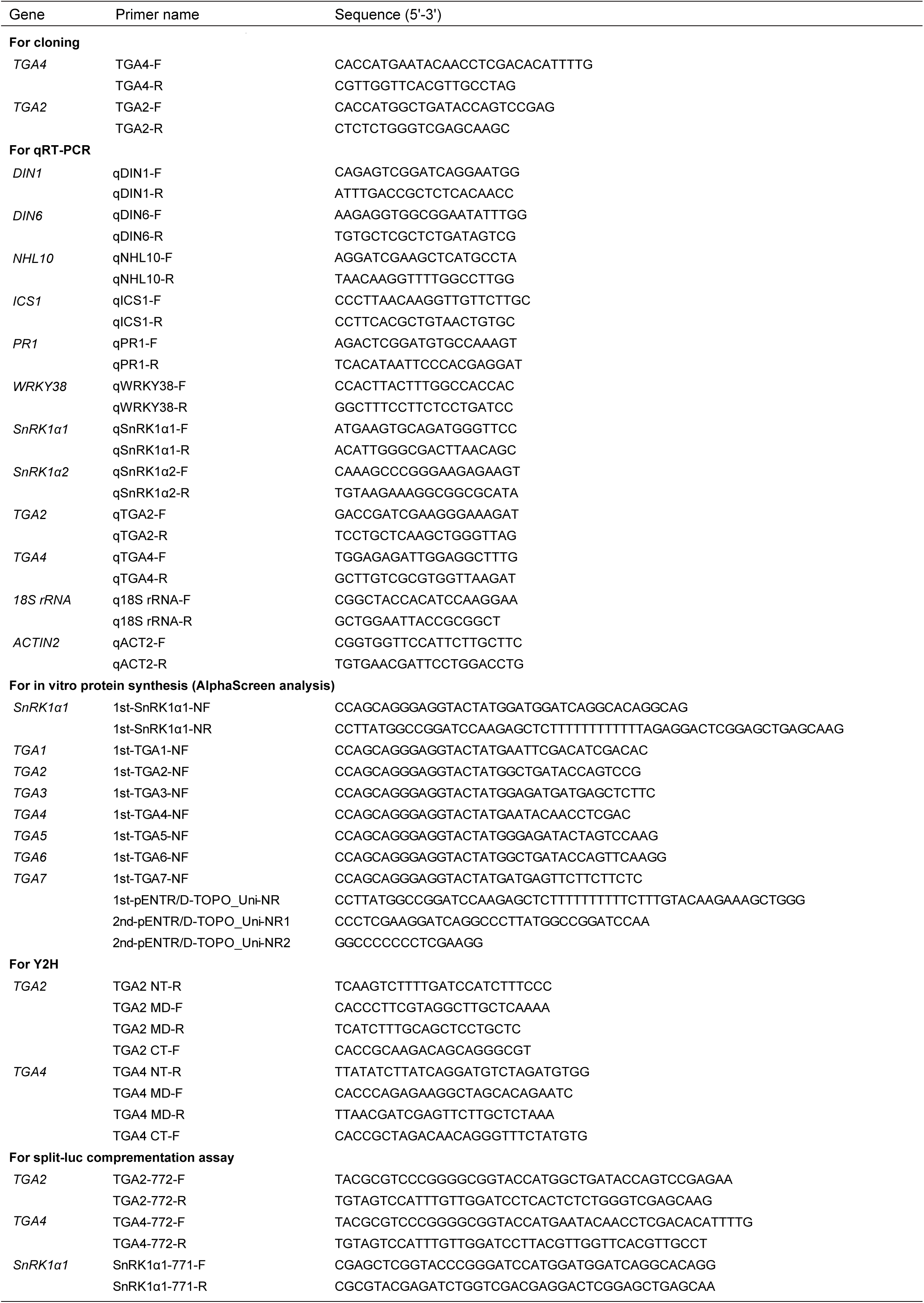
List of primers used in this study.

## Notes

### Competing Interest Statement

The authors have declared no competing interest.

## References

1. R. B. Abramovitch, J. C. Anderson, G. B. Martin, Bacterial elicitation and evasion of plant innate immunity. Nat. Rev. Mol. Cell Biol. 7, 601–611 (2006).

2. J. D. G. Jones, J. L. Dangl, The plant immune system. Nature 444, 323–329 (2006).

3. S. T. Chisholm, G. Coaker, B. Day, B. J. Staskawicz, Host-Microbe Interactions: Shaping the Evolution of the Plant Immune Response. Cell 124, 803–814 (2006).

4. D. Couto, C. Zipfel, Regulation of pattern recognition receptor signalling in plants. Nat. Rev. Immunol. 16, 537–552 (2016).

5. M. Bjornson, P. Pimprikar, T. Nürnberger, C. Zipfel, The transcriptional landscape of Arabidopsis thaliana pattern-triggered immunity. Nat. Plants 7, 579–586 (2021).

6. M. Yuan, B. P. M. Ngou, P. Ding, X.-F. Xin, PTI-ETI crosstalk: an integrative view of plant immunity. Curr. Opin. Plant Biol. 62, 102030 (2021).

7. C. An, Z. Mou, Salicylic Acid and its Function in Plant Immunity. J. Integr. Plant Biol. 53, 412–428 (2011).

8. P. Ding, Y. Ding, Stories of Salicylic Acid: A Plant Defense Hormone. Trends Plant Sci. 25, 549–565 (2020).

9. Z. Q. Fu, X. Dong, Systemic Acquired Resistance: Turning Local Infection into Global Defense. Annu. Rev. Plant Biol. 64, 839–863 (2013).

10. Y. Peng, J. Yang, X. Li, Y. Zhang, Salicylic Acid: Biosynthesis and Signaling. Annu. Rev. Plant Biol. 72, 761–791 (2021).

11. M. C. Wildermuth, J. Dewdney, G. Wu, F. M. Ausubel, Isochorismate synthase is required to synthesize salicylic acid for plant defence. Nature 414, 562–565 (2001).

12. M. P. Torrens-Spence, et al., PBS3 and EPS1 Complete Salicylic Acid Biosynthesis from Isochorismate in Arabidopsis. Mol. Plant 12, 1577–1586 (2019).

13. Y. Wu, et al., The Arabidopsis NPR1 Protein Is a Receptor for the Plant Defense Hormone Salicylic Acid. Cell Rep. 1, 639–647 (2012).

14. Y. Ding, et al., Opposite Roles of Salicylic Acid Receptors NPR1 and NPR3/NPR4 in Transcriptional Regulation of Plant Immunity. Cell 173, 1454–1467.e15 (2018).

15. M. Kesarwani, J. Yoo, X. Dong, Genetic Interactions of TGA Transcription Factors in the Regulation of Pathogenesis-Related Genes and Disease Resistance in Arabidopsis. Plant Physiol. 144, 336–346 (2007).

16. C. Després, C. DeLong, S. Glaze, E. Liu, P. R. Fobert, The Arabidopsis NPR1/NIM1 Protein Enhances the DNA Binding Activity of a Subgroup of the TGA Family of bZIP Transcription Factors. Plant Cell 12, 279–290 (2000).

17. M. Jakoby, et al., bZIP transcription factors in Arabidopsis. Trends Plant Sci. 7, 106–111 (2002).

18. C. Xiang, Z. Miao, E. Lam, DNA-binding properties, genomic organization and expression pattern of TGA6, a new member of the TGA family of bZIP transcription factors in Arabidopsis thaliana. Plant Mol. Biol. 34, 403–415 (1997).

19. C. Després, et al., The Arabidopsis NPR1 Disease Resistance Protein Is a Novel Cofactor That Confers Redox Regulation of DNA Binding Activity to the Basic Domain/Leucine Zipper Transcription Factor TGA1. Plant Cell 15, 2181–2191 (2003).

20. J.-M. Zhou, et al., NPR1 Differentially Interacts with Members of the TGA/OBF Family of Transcription Factors That Bind an Element of the PR-1 Gene Required for Induction by Salicylic Acid. MPMI 13, 191–202 (2000).

21. C. Johnson, E. Boden, J. Arias, Salicylic Acid and NPR1 Induce the Recruitment of *trans-*Activating TGA Factors to a Defense Gene Promoter in Arabidopsis. Plant Cell 15, 1846–1858 (2003).

22. A. Rochon, P. Boyle, T. Wignes, P. R. Fobert, C. Després, The Coactivator Function of *Arabidopsis* NPR1 Requires the Core of Its BTB/POZ Domain and the Oxidation of C-Terminal Cysteines. Plant Cell 18, 3670–3685 (2007).

23. H. Jin, et al., Salicylic acid-induced transcriptional reprogramming by the HAC– NPR1–TGA histone acetyltransferase complex in Arabidopsis. Nucleic Acids Res. 46, 11712–11725 (2018).

24. Y. Zhang, M. J. Tessaro, M. Lassner, X. Li, Knockout Analysis of Arabidopsis Transcription Factors *TGA2*, *TGA5*, and *TGA6* Reveals Their Redundant and Essential Roles in Systemic Acquired Resistance. Plant Cell 15, 2647–2653 (2003).

25. S. Kumar, et al., Structural basis of NPR1 in activating plant immunity. Nature 605, 561–566 (2022).

26. Y.-L. Chen, et al., XCP1 cleaves Pathogenesis-related protein 1 into CAPE9 for systemic immunity in Arabidopsis. Nat. Commun. 14, 4697 (2023).

27. K.-C. Kim, Z. Lai, B. Fan, Z. Chen, *Arabidopsis* WRKY38 and WRKY62 Transcription Factors Interact with Histone Deacetylase 19 in Basal Defense. Plant Cell 20, 2357–2371 (2008).

28. L. Yao, et al., High air humidity dampens salicylic acid pathway and NPR1 function to promote plant disease. EMBO J. 42, e113499 (2023).

29. J. H. Kim, et al., Increasing the resilience of plant immunity to a warming climate. Nature 607, 339–344 (2022).

30. S. Li, et al., INDUCER OF CBF EXPRESSION 1 promotes cold-enhanced immunity by directly activating salicylic acid signaling. Plant Cell 36, 2587–2606 (2024).

31. J. Sheen, L. Zhou, J.-C. Jang, Sugars as signaling molecules. Curr. Opin. Plant Biol. 2, 410–418 (1999).

32. K. Koch, Sucrose metabolism: regulatory mechanisms and pivotal roles in sugar sensing and plant development. Curr. Opin. Plant Biol. 7, 235–246 (2004).

33. F. Rolland, E. Baena-Gonzalez, J. Sheen, SUGAR SENSING AND SIGNALING IN PLANTS: Conserved and Novel Mechanisms. Annu. Rev. Plant Biol. 57, 675–709 (2006).

34. M. R. Bolouri-Moghaddam, K. Le Roy, L. Xiang, F. Rolland, W. Van Den Ende, Sugar signalling and antioxidant network connections in plant cells. FEBS J. 277, 2022–2037 (2010).

35. B. Moore, et al., Role of the *Arabidopsis* Glucose Sensor HXK1 in Nutrient, Light, and Hormonal Signaling. Science 300, 332–336 (2003).

36. T. Broeckx, S. Hulsmans, F. Rolland, The plant energy sensor: evolutionary conservation and divergence of SnRK1 structure, regulation, and function. J. Exp. Bot. 67, 6215–6252 (2016).

37. L. Shi, Y. Wu, J. Sheen, TOR signaling in plants: conservation and innovation. Development 145, dev160887 (2018).

38. T. Dobrenel, et al., TOR Signaling and Nutrient Sensing. Annu. Rev. Plant Biol. 67, 261–285 (2016).

39. Y. Xiong, et al., Glucose–TOR signalling reprograms the transcriptome and activates meristems. Nature 496, 181–186 (2013).

40. E. Baena-González, F. Rolland, J. M. Thevelein, J. Sheen, A central integrator of transcription networks in plant stress and energy signalling. Nature 448, 938–942 (2007).

41. Y.-H. Cho, J.-W. Hong, E.-C. Kim, S.-D. Yoo, Regulatory Functions of SnRK1 in Stress-Responsive Gene Expression and in Plant Growth and Development. Plant Physiol. 158, 1955–1964 (2012).

42. H.-Y. Cho, E. Loreti, M.-C. Shih, P. Perata, Energy and sugar signaling during hypoxia. New Phytol. 229, 57–63 (2021).

43. A. Mair, et al., SnRK1-triggered switch of bZIP63 dimerization mediates the low-energy response in plants. eLife 4, e05828 (2015).

44. C. Weiste, et al., The Arabidopsis bZIP11 transcription factor links low-energy signalling to auxin-mediated control of primary root growth. PLOS Genet. 13, e1006607 (2017).

45. Y. Fujiki, et al., Dark-inducible genes from *Arabidopsis thaliana* are associated with leaf senescence and repressed by sugars. Physiol. Plant. 111, 345–352 (2001).

46. H.-Y. Cho, M.-Y. J. Lu, M.-C. Shih, The SnRK1-eIFiso4G1 signaling relay regulates the translation of specific mRNAs in Arabidopsis under submergence. New Phytol. 222, 366–381 (2019).

47. H.-Y. Cho, T.-N. Wen, Y.-T. Wang, M.-C. Shih, Quantitative phosphoproteomics of protein kinase SnRK1 regulated protein phosphorylation in Arabidopsis under submergence. J. Exp. Bot. 67, 2745–2760 (2016).

48. C. Polge, M. Thomas, SNF1/AMPK/SnRK1 kinases, global regulators at the heart of energy control? Trends Plant Sci. 12, 20–28 (2007).

49. S. Emanuelle, M. S. Doblin, D. I. Stapleton, A. Bacic, P. R. Gooley, Molecular Insights into the Enigmatic Metabolic Regulator, SnRK1. Trends Plant Sci. 21, 341–353 (2016).

50. R. Ghillebert, et al., The AMPK/SNF1/SnRK1 fuel gauge and energy regulator: structure, function and regulation. FEBS J. 278, 3978–3990 (2011).

51. J. M. Gancedo, Yeast Carbon Catabolite Repression. Microbiol. Mol. Biol. Rev. 62, 334–361 (1998).

52. M. Carlson, Glucose repression in yeast. Curr. Opin. Microbiol. 2, 202–207 (1999).

53. C.-S. Zhang, et al., Fructose-1,6-bisphosphate and aldolase mediate glucose sensing by AMPK. Nature 548, 112–116 (2017).

54. Y. Zong, et al., Hierarchical activation of compartmentalized pools of AMPK depends on severity of nutrient or energy stress. Cell Res. 29, 460–473 (2019).

55. Z. Zhai, et al., Trehalose 6-Phosphate Positively Regulates Fatty Acid Synthesis by Stabilizing WRINKLED1. Plant Cell 30, 2616–2627 (2018).

56. J. Van Leene, et al., Mapping of the plant SnRK1 kinase signalling network reveals a key regulatory role for the class II T6P synthase-like proteins. Nat. Plants 8, 1245–1261 (2022).

57. M. Sanagi, et al., Low nitrogen conditions accelerate flowering by modulating the phosphorylation state of FLOWERING BHLH 4 in *Arabidopsis*. Proc. Natl. Acad. Sci. 118, e2022942118 (2021).

58. P. Muralidhara, et al., Perturbations in plant energy homeostasis prime lateral root initiation via SnRK1-bZIP63-ARF19 signaling. Proc. Natl. Acad. Sci. 118, e2106961118 (2021).

59. R. Szczesny, et al., Suppression of the AvrBs1-specific hypersensitive response by the YopJ effector homolog AvrBsT from *Xanthomonas* depends on a SNF1-related kinase. New Phytol. 187, 1058–1074 (2010).

60. O. Filipe, D. De Vleesschauwer, A. Haeck, K. Demeestere, M. Höfte, The energy sensor OsSnRK1a confers broad-spectrum disease resistance in rice. Sci. Rep. 8, 3864 (2018).

61. W. Shen, et al., Sucrose Nonfermenting 1-Related Protein Kinase 1 Phosphorylates a Geminivirus Rep Protein to Impair Viral Replication and Infection. Plant Physiol. 178, 372–389 (2018).

62. R. Ehness, M. Ecker, D. E. Godt, T. Roitsch, Glucose and Stress Independently Regulate Source and Sink Metabolism and Defense Mechanisms via Signal Transduction Pathways Involving Protein Phosphorylation. Plant Cell 9, 1825–1841 (1997).

63. J. Gómez-Ariza, et al., Sucrose-Mediated Priming of Plant Defense Responses and Broad-Spectrum Disease Resistance by Overexpression of the Maize Pathogenesis-Related PRms Protein in Rice Plants. MPMI 20, 832–842 (2007).

64. T. Engelsdorf, et al., Reduced Carbohydrate Availability Enhances the Susceptibility of Arabidopsis toward *Colletotrichum higginsianum*. Plant Physiol. 162, 225–238 (2013).

65. K. Yamada, A. Mine, Sugar coordinates plant defense signaling. Sci. Adv. 10, eadk4131 (2024).

66. E. Baena-González, F. Rolland, J. M. Thevelein, J. Sheen, A central integrator of transcription networks in plant stress and energy signalling. Nature 448, 938–942 (2007).

67. M. Boudsocq, et al., Differential innate immune signalling via Ca^2+^ sensor protein kinases. Nature 464, 418–422 (2010).

68. S. Uknes, B. Mauch-Mani, D. Chandler, A. Slusarenko, Acquired Resistance in Arabidopsis. Plant Cell 4, 645–656 (1992).

69. J. Dong, C. Chen, Z. Chen, Expression profiles of the Arabidopsis WRKY gene superfamily during plant defense response. Plant Mol. Biol. 51, 21–37 (2003).

70. M. Sanagi, et al., Sugar-responsive transcription factor bZIP3 affects leaf shape in Arabidopsis plants. Plant Biotechnol. 35, 167–170 (2018).

71. S. Huang, et al., Identification and receptor mechanism of TIR-catalyzed small molecules in plant immunity. Science 377, eabq3297 (2022).

72. A. Jia, et al., TIR-catalyzed ADP-ribosylation reactions produce signaling molecules for plant immunity. Science 377, eabq8180 (2022).

73. M. Hartmann, et al., Flavin Monooxygenase-Generated N-Hydroxypipecolic Acid Is a Critical Element of Plant Systemic Immunity. Cell 173, 456–469.e16 (2018).

74. H. Cao, S. A. Bowling, A. S. Gordon, Characterization of an Arabidopsis Mutant That Is Nonresponsive to Inducers of Systemic Acquired Resistance. Plant Cell 6, 1583–1592 (1994).

75. Y. T. Cheng, L. Zhang, S. Y. He, Plant-Microbe Interactions Facing Environmental Challenge. Cell Host Microbe 26, 183–192 (2019).

76. B. K. Singh, et al., Climate change impacts on plant pathogens, food security and paths forward. Nat. Rev. Microbiol. 21, 640–656 (2023).

77. C. Roussin-Léveillée, C. A. M. Rossi, C. D. M. Castroverde, P. Moffett, The plant disease triangle facing climate change: a molecular perspective. Trends Plant Sci. 29, 895–914 (2024).

78. M. Nomoto, Y. Tada, Cloning-free template DNA preparation for cell-free protein synthesis via two-step PCR using versatile primer designs with short 3′-UTR. Genes Cells 23, 46–53 (2018).

79. M. Nomoto, et al., Suppression of MYC transcription activators by the immune cofactor NPR1 fine-tunes plant immune responses. Cell Rep. 37, 110125 (2021).

80. G. Zhou, et al., Role of AMP-activated protein kinase in mechanism of metformin action. J. Clin. Invest. 108, 1167–1174 (2001).

81. S. Yuan, et al., *Arabidopsis* cryptochrome 1 functions in nitrogen regulation of flowering. Proc. Natl. Acad. Sci. 113, 7661–7666 (2016).

82. R. B. Stevens, “Cultural Practices in Disease Control” in Plant Pathology, An Advanced Treatise, ed. J. G. Horsfall, A. E. Diamond (Academic Press), pp, 357–429 (1960).

83. B. Huot, et al., Dual impact of elevated temperature on plant defence and bacterial virulence in Arabidopsis. Nat. Commun. 8, 1808 (2017).

84. Y. Zhu, W. Qian, J. Hua, Temperature Modulates Plant Defense Responses through NB-LRR Proteins. PLoS Pathog. 6, e1000844 (2010).

85. S. Bhattacharjee, M. K. Halane, S. H. Kim, W. Gassmann, Pathogen Effectors Target *Arabidopsis* EDS1 and Alter Its Interactions with Immune Regulators. Science 334, 1405–1408 (2011).

86. K. Heidrich, et al., Arabidopsis TNL-WRKY domain receptor RRS1 contributes to temperature-conditioned RPS4 auto-immunity. Front. Plant Sci. 4, 403 (2013).

87. H. Demont, et al., Downstream signaling induced by several plant Toll/interleukin-1 receptor-containing immune proteins is stable at elevated temperature. Cell Rep. 44, 115326 (2025).

88. X.-F. Xin, et al., Bacteria establish an aqueous living space in plants crucial for virulence. Nature 539, 524–529 (2016).

89. Y. Hu, et al., Bacterial effectors manipulate plant abscisic acid signaling for creation of an aqueous apoplast. Cell Host Microbe 30, 518–529.e6 (2022).

90. C. Roussin-Léveillée, et al., Evolutionarily conserved bacterial effectors hijack abscisic acid signaling to induce an aqueous environment in the apoplast. Cell Host Microbe 30, 489–501.e4 (2022).

91. S. Yasuda, et al., Humidity-driven ABA depletion determines plant-pathogen competition for leaf water. bioRxiv (2025) doi: 10.1101/2025.01.04.631318.

92. A. Rodrigues, et al., ABI1 and PP2CA Phosphatases Are Negative Regulators of Snf1-Related Protein Kinase1 Signaling in *Arabidopsis*. Plant Cell 25, 3871–3884 (2013).

93. B. Belda-Palazón, et al., A dual function of SnRK2 kinases in the regulation of SnRK1 and plant growth. Nat. Plants 6, 1345–1353 (2020).

94. G. Qi, et al., Pandemonium Breaks Out: Disruption of Salicylic Acid-Mediated Defense by Plant Pathogens. Mol. Plant 11, 1427–1439 (2018).

95. T. Sun, et al., TGACG-BINDING FACTOR 1 (TGA1) and TGA4 regulate salicylic acid and pipecolic acid biosynthesis by modulating the expression of *SYSTEMIC ACQUIRED RESISTANCE DEFICIENT 1* (*SARD1*) and *CALMODULIN-BINDING PROTEIN 60g* (*CBP60g*). New Phytol. 217, 344–354 (2018).

96. Y.-W. Kim, et al., Brassinosteroids enhance salicylic acid-mediated immune responses by inhibiting BIN2 phosphorylation of clade I TGA transcription factors in Arabidopsis. Mol. Plant 15, 991–1007 (2022).

97. Q. Han, et al., Salicylic acid-activated BIN2 phosphorylation of TGA3 promotes *Arabidopsis* PR gene expression and disease resistance. EMBO J. 41, e110682 (2022).

## SI References

1. M. Sanagi, et al., Low nitrogen conditions accelerate flowering by modulating the phosphorylation state of FLOWERING BHLH 4 in *Arabidopsis*. Proc. Natl. Acad. Sci. 118, e2022942118 (2021).

2. M. Sanagi, et al., Sugar-responsive transcription factor bZIP3 affects leaf shape in Arabidopsis plants. Plant Biotechnol. 35, 167–170 (2018).

3. M. C. Wildermuth, J. Dewdney, G. Wu, F. M. Ausubel, Isochorismate synthase is required to synthesize salicylic acid for plant defence. Nature 414, 562–565 (2001).

4. H. Cao, S. A. Bowling, A. S. Gordon, Characterization of an Arabidopsis Mutant That 1s Nonresponsive to lnducers of Systemic Acquired Resistance. Plant Cell 6, 1583–1592 (1994)

5. M. Kesarwani, J. Yoo, X. Dong, Genetic Interactions of TGA Transcription Factors in the Regulation of Pathogenesis-Related Genes and Disease Resistance in Arabidopsis. Plant Physiol. 144, 336–346 (2007).

6. Y. Zhang, M. J. Tessaro, M. Lassner, X. Li, Knockout Analysis of Arabidopsis Transcription Factors *TGA2*, *TGA5*, and *TGA6* Reveals Their Redundant and Essential Roles in Systemic Acquired Resistance. Plant Cell 15, 2647–2653 (2003).

7. K. Yamada, A. Mine, Sugar coordinates plant defense signaling. Sci. Adv. 10, eadk4131 (2024).

8. T. Nakagawa, et al., Development of series of gateway binary vectors, pGWBs, for realizing efficient construction of fusion genes for plant transformation. J. Biosci. Bioeng. 104, 34–41 (2007).

9. S. Chen, Y. Zhou, Y. Chen, J. Gu, fastp: an ultra-fast all-in-one FASTQ preprocessor. Bioinformatics 34, i884–i890 (2018).

10. A. Dobin, et al., STAR: ultrafast universal RNA-seq aligner. Bioinformatics 29, 15–21 (2013).

11. Y. Liao, G. K. Smyth, W. Shi, featureCounts: an efficient general purpose program for assigning sequence reads to genomic features. Bioinformatics 30, 923–930 (2014).

12. M. I. Love, W. Huber, S. Anders, Moderated estimation of fold change and dispersion for RNA-seq data with DESeq2. Genome Biol. 15, 550 (2014).

13. T. Wu, et al., clusterProfiler 4.0: A universal enrichment tool for interpreting omics data. The Innovation 2, 100141 (2021).

14. R. Vera Alvarez, L. S. Pongor, L. Mariño-Ramírez, D. Landsman, TPMCalculator: one-step software to quantify mRNA abundance of genomic features, Bioinformatics 35, 1960–1962 (2019)

15. Y. Tsunoda, et al., Improving expression and solubility of rice proteins produced as fusion proteins in Escherichia coli. Protein Expr. Purif. 42, 268–277 (2005).

16. M. Nomoto, Y. Tada, Cloning-free template DNA preparation for cell-free protein synthesis via two-step PCR using versatile primer designs with short 3′-UTR. Genes Cells 23, 46–53 (2018).

17. M. Nomoto, et al., Suppression of MYC transcription activators by the immune cofactor NPR1 fine-tunes plant immune responses. Cell Rep. 37, 110125 (2021).

18. Q. Lu, et al., Arabidopsis homolog of the yeast TREX-2 mRNA export complex: components and anchoring nucleoporin: TREX-2 mRNA export complex. Plant J. 61, 259–270 (2009).

19. J.-Y. Gou, et al., Negative Regulation of Anthocyanin Biosynthesis in Arabidopsis by a miR156-Targeted SPL Transcription Factor, Plant Cell 23, 1512–1522 (2011).

